# Mapping the nervous system of the Idiosepius *hallami* pygmy squid: insights from whole-animal X-ray nanotomography imaging

**DOI:** 10.1101/2025.09.25.678516

**Authors:** Ana Correia, Wen-Sung Chung, Samia Mohinta, Daniel Franco-Barranco, Artem Vorobyev, Alexandra Pacureanu, Albert Cardona, Marc Corrales

**Affiliations:** MRC Laboratory of Molecular Biology, Cambridge, UK; Department of Physiology, Development and Neuroscience, University of Cambridge, Cambridge, UK; School of the Environment, The University of Queensland, St Lucia 4072, QLD, Australia; Donostia International Physics Center (DIPC), San Sebastian, Spain; ESRF, The European Synchrotron, Grenoble, France

## Abstract

The study of a nervous system as big as the cephalopod’s requires multimodal imaging approaches capable of capturing neural architecture across scales. Here, we present a whole-animal volume of the pygmy squid hatchling *Idiosepius hallami*, acquired using X-ray holographic nanotomography at the beamline ID16A of the European Synchrotron. The reconstructed 3D volume comprises 40 tiled scans acquired at a voxel size of 125 nm. While individual neurons are not resolved at this resolution, we segmented major body regions and mapped the large-scale connectivity by tracing afferent and efferent nerve bundles, including projections from the olfactory organs, chromatophore lobes, and arm ganglia to the brain. The acquisition of this dataset represents a significant milestone for X-ray nanotomography, being the largest whole animal volume imaged at this spatial resolution. The volume serves as a resource for comparative neuroscience and cephalopod biology.

## Introduction

Cephalopods evolved from monoplacophoran-like molluscs during the Cambrian explosion over 540 million years ago (Strugnell et al., 2006). Approximately 800 different species of extant cephalopods inhabit marine or brackish environments (Jereb and Roper, 2005, 2010; Jereb et al., 2016), with body sizes ranging from about 10 millimetres (e.g., adult *Idiosepius* pygmy squid) to 13 meters (e.g., *Mesonychoteuthis hamiltoni,* the colossal squid). The coleoid cephalopods (octopus, squid, and cuttlefish, but not nautilus) are characterised by their streamlined body shape, advanced sensory systems and centralised multi-lobed brain (Wells and Wells, 1957; Nixon and Young, 2003; Chung and Marshall, 2014; Marini et al., 2017; Shigeno et al., 2018; Chung, Kurniawan and Marshall, 2020, 2022; Chung et al., 2023; Pungor and Niell, 2023). Coleoids have a short life span (1-2 years) and face varying challenges throughout their lives, including changing environments, different predator-prey interactions, and a bet-hedging semelparous reproductive strategy (Roger T. Hanlon and Messenger, 2018). These changes in ecological and environmental demands are met by an elaborate cognitive flexibility, which is based on their large and complex brain as they learn and mature (Nixon and Young, 2003). While the overall Bauplan appears distinct, there are similarities between the brains of coleoids and vertebrates, particularly in neurogenesis during development and function (Shigeno et al., 2018; Deryckere et al., 2021; Chung, Kurniawan and Marshall, 2022). For example, the optic lobe (OPL) equates to the tectum and the inferior frontal lobe complex (iFLx) to the olfactory bulb. These soft-bodied animals also exhibit a striking and unique repertoire of complex behaviours, including colour-blind camouflage (Marshall and Messenger, 1996; Chung and Marshall, 2016), courtship display (Lin, Tsai and Chiao, 2017), mate guarding (Huffard, Caldwell and Boneka, 2008), interspecific collaborative hunting (Sampaio et al., 2024) and observational learning (Fiorito and Scotto, 1992). Their complex and large brains rapidly integrate sensory information and coordinate nuanced motor control that underpin their advanced cognitive activities (Young, 1961; Boycott, 1965; Fiorito, Planta and Scotto, 1990; Messenger, 2001; How et al., 2017; Turchetti-Maia, Shomrat and Hochner, 2017; Reiter et al., 2018; Roger T Hanlon and Messenger, 2018; Shigeno et al., 2018; Amodio et al., 2019; Schnell and Clayton, 2019; Osorio et al., 2022).

Like other molluscs, the cephalopod central nervous system (CNS) comprises central ganglia or medullary nerves surrounding the oesophagus (Yamamoto, Shimazaki and Shigeno, 2003; Hochner and Glanzman, 2016; Koizumi et al., 2016; Chung, Kurniawan and Marshall, 2020; Chung et al., 2023; Montague et al., 2023). Distinct brain lobes each feature a central neuropil surrounded by a perikaryal layer. The brain is divided in four main parts: the subesophageal mass (motor control, e.g. arm and fin movement); the supraesophageal mass (a higher motor centre integrating sensory inputs with motor output); the vertical complex (learning and memory); and two optic lobes (visual information processing) (Wollesen, 2015; Chung, Kurniawan and Marshall, 2020; Chung et al., 2023; Montague et al., 2023; Pungor et al., 2023).

The pioneering neuroanatomical research of Cajal and J. Z. Young extensively describes cephalopod cell types and possible synapses between neurons using only Golgi stainings and single-section Transmission Electron Microscopy (TEM) (Cajal, 1917; Young, 1961, 1962, 1971, 1974a, 1974b, 1976, 1977, 1979). These foundational studies, combined with brain lobe ablation experiments mapping structure to behaviour (Boycott and Young, 1957; Wells and Wells, 1957; Boycott, 1961, 1965; Abbott, Williamson and Maddock, 1995), established cephalopods as a model organism early on in research on nervous system development (Ponte et al., 2021; Styfhals et al., 2022; Albertin and Katz, 2023), learning (Boycott, 1965; Fiorito, Planta and Scotto, 1990; Marini et al., 2017; Schnell et al., 2021; Jozet-Alves, Schnell and Clayton, 2023), motor control (Kuuspalu, Cody and Hale, 2022), vision (Chung and Marshall, 2014; Pungor and Niell, 2023; Pungor et al., 2023), camouflage (Hanlon, 2007; Reiter et al., 2018; Osorio et al., 2022; Woo et al., 2023) and behaviour (Roger T Hanlon and Messenger, 2018). Atlases, in which brains are decomposed in lobes, and projectomes describing their major connections, abound in the literature for different cephalopod species (Yamamoto, Shimazaki and Shigeno, 2003; Koizumi et al., 2016; Styfhals et al., 2022; Chung et al., 2023; Montague et al., 2023).

The pygmy squid (family Idiosepiidae) is a group of small-sized squids dwelling in the tropical Indo-Pacific coastal waters (Nishiguchi et al., 2014; Reid and Strugnell, 2018). One unique feature of the pygmy squid is that its adhesive gland secretes sticky mucus that allows the squid to attach itself to substrates (e.g. seagrass or coral rubble) (Byern et al., 2008; Nishiguchi et al., 2014). They are sexually dimorphic: males reach ∼15 mm, females ∼21 mm in mantle length (Nishiguchi et al., 2014; Reid and Strugnell, 2018). Despite being the smallest cephalopod, recent studies showed that pygmy squids exhibit complex behaviours similar to those of their larger-sized siblings, including colour display during courtship, and use of cephalotoxins and ink during predation (Shigeno and Yamamoto, 2002; Sato et al., 2016; Reid and Strugnell, 2018). Both sexes reach sexual maturity within two months post-hatching. In contrast to most cephalopods which are semelparous, the female pygmy squid repeatedly lays small clutches of eggs across weeks (30-80 every 2-7 days for a month) and their eggs take approximately 2-3 weeks to hatch (*I. hallami*, at 25oC, Chung et al. unpublished), making them a promising animal model to study neurobiology in the lab and to map their whole-brain connectome (Shigeno and Yamamoto, 2002; Kasugai and Segawa, 2005; Nishiguchi et al., 2014).

Advances in synchrotron-based X-ray imaging, paired with suitable staining protocols for whole-animal and whole-brain samples (Mikula and Denk, 2015; Genoud et al., 2018; Lu et al., 2022, 2023; Ströh et al., 2022; Song, Feng and Helmstaedter, 2023), make multimodal connectomics possible. Light sources such as the European Synchrotron (ESRF) produce X-ray beams 10 trillion times brighter than standard hospital X-ray instruments. The high photon flux is combined with spatial and temporal coherence to leverage phase contrast and drastically improve the signal-to-noise ratio. Specifically, the ESRF beamline ID16A (used in this study) delivers images at 20 to 130 nm per voxel (Silva et al., 2017; Villar et al., 2018). A recent example is Kuan et al. (2020), who acquired a dataset with sub-100 nm resolution over millimetre-scale volumes with X-ray holographic nanotomography (XNH) at the ESRF ID16A beamline, later correlating it with EM (Kuan et al. 2020a).

To bridge the scales needed for a faithful description of the cephalopod nervous system, we developed a sample preparation protocol that supports a multi-modal imaging approach, which can sequentially combine synchrotron X-ray holographic nanotomography and electron microscopy, in addition to carefully selecting our model organism, the Australian pygmy squid *Idiosepius hallami*, of suitably reduced dimensions and readily available. Then, we imaged a volume of a whole pygmy squid hatchling using the ESRF beamline ID16A at 125 nm isotropic resolution, and segmented different structures, including the brain lobes, sensory systems and the associated nerve tracts.

## Methods

### Fixation

All collections were conducted under the Queensland General Fisheries Permit (207976) and Marine Park Permits (QS2013/CVL625, MPP19-002266). The sexually mature pygmy squid, Idiosepius hallami, were collected using hand nets (water depth 0-0.2 m during the low tide) from the seagrass meadow close to Moreton Bay Research Station (MBRS; 27.4966° S, 153.4000° E), Minjerribah, Queensland, Australia. The animals were transferred to the MBRS aquarium facility, where the selected pairs were kept in the holding tank (88 cm x 44 cm x 31 cm). Several green-coloured plastic thin strips (15cm x 1 cm) were placed in the holding tank to allow females to lay eggs. The egg clutches were then transferred in the brooding boxes (15 cm x 10 cm x 7 cm) floating in the holding tank. The maintenance and experimental protocol were approved by the University of Queensland’s Animal Ethics Committee NEWMA (2016/AE000304, 2022/AE000389). I. hallami hatchlings (30 minutes post-hatching) were anaesthetised in seawater mixed with 2% MgCl2 (Chem-Supply, Australia) and sacrificed by an overdose of MgCl2 prior to fixation. The samples were fixed with 2.5% glutaraldehyde, 2% paraformaldehyde (EM grade, Electron Microscopy Sciences, Hatfield, USA) in 0.15M sodium cacodylate (pH 7.4) at 4 ℃ at the University of Queensland, Australia, and shipped in ice packs to the Laboratory of Molecular Biology. We estimate the time of fixation to be ∼72h. Upon arrival, samples were washed for 30 min in 0.15M sodium cacodylate, and the protocol continued as detailed in Table 1. The samples were stained with osmium tetroxide, enabling multimodal imaging, using both electron microscopy and X-ray nanoholotomography techniques. In brief, the BROPA/HUA protocol, developed for the whole mouse brain, was adapted to fix the entire animal, with the primary purpose of minimising cracks in the tissue, and having it completely stained throughout, without the presence of a staining gradient.

**Table 1.** Protocol for synchrotron X-ray imaging. OSO4 - Osmium Tetroxide, K4Fe(CN)6 - Potassium Ferrocyanide, UA - Uranyl Acetate

| Step | Solution | Time | Temperature | Agitation | Dark |
| --- | --- | --- | --- | --- | --- |
| Fixation | 2.5% glutaraldehyde, 2% paraformaldehyde in 0.15M sodium cacodylate (pH 7.4) | 72hr | 4°C | Yes | No |
| Washes | 0.15M sodium cacodylate (pH 7.4) | 1x30min | RT | Yes | No |
| Osmication | 2% OSO<sub>4</sub> in buffer | >72hr | 4°C | Yes | Yes |
| washes | 0.15M sodium cacodylate (pH 7.4) | 3X30min | RT | Yes | No |
| Reduction | 2.5% K<sub>4</sub>Fe(CN)<sub>6</sub> in buffer | O/N | 4°C | Yes | Yes |
| Washes | 0.15M sodium cacodylate (pH 7.4) | 3X30min | RT | Yes | No |
| Osmication | 2% OSO<sub>4</sub> in buffer | 3hr | 4°C | Yes | Yes |
| Washes | 0.15M sodium cacodylate (pH 7.4) | 2X30min | RT | Yes | No |
| Washes | ddH<sub>2</sub>O | 2X30min | RT | Yes | No |
| Pyrogallol | 4% Pyrogallol in ddH<sub>2</sub>O | O/N | RT | Yes | Yes |
| Washes | ddH<sub>2</sub>O | 2X30min | RT | Yes | No |
| Osmication | 2% OSO<sub>4</sub> in ddH<sub>2</sub>O | 6hr | ICE | Yes | Yes |
| Washes | ddH<sub>2</sub>O | 2X30min | RT | Yes | No |
| Bulk Stain | 2% UA in ddH<sub>2</sub>O | O/N | 4°C | Yes | Yes |
| Bulk Stain | 2% UA in ddH<sub>2</sub>O | 2hr | 50°C | Yes | Yes |
| Washes | ddH<sub>2</sub>O | 3X10min | RT | Yes | No |
| Lead Staining | Lead aspartate (pH 5.5 -5.8)<br>Lead aspartate (pH 5.5 -5.8) | 30min<br>2hr | 65°C<br>RT | No<br>Yes | Yes<br>Yes |
| Washes | ddH<sub>2</sub>O | 5X5min | RT | Yes | No |
| Dehydration | 10, 30, 50, 70, 80, 90 and 95% acetone<br>100% acetone<br>100% acetone | 10 min each step<br>2X10 min<br>2X10 min | ICE<br>ICE<br>RT | Yes<br>Yes<br>Yes | No<br>No<br>No |
| Infiltration in resin: durcupan or embed812 | (1:2) Resin in acetone<br>(1:1) Resin in acetone<br>(2:1) Resin in acetone<br>100% Resin | 5hr<br>12hr<br>5hr<br>12hr | RT<br>RT<br>RT<br>RT | Yes<br>Yes<br>Yes<br>Yes | No<br>No<br>No<br>No |
| Polymerisation | Group A: Fresh 100% Resin<br>Group A: 100% Resin in embedding mould | 4-6hr<br>48hr | RT<br>60°C | Yes<br>No | No<br>Yes |

Given the different resistance of different resin formulations to X-rays, samples were infiltrated and embedded with Durcupan resin (44610 Sigma-Aldrich) or Embed812 resin (14120 EMS). Both resins were cured for 48 hours at 70℃.

### Sample selection and mounting

All cured blocks were scanned using a Bruker Skyscan 2214 microCT system. The microCT screening enabled a quality control assessment, allowing us to verify the presence of artefacts such as cracks in the tissue and evaluate the staining throughout the pygmy squid. Abundant heavy metal impregnation, while optimal for electron microscopy imaging and able to provide contrast to X-rays, results in pronounced X-ray absorption. This phenomenon increases sample heating and warping. If any cracks are present, they can expand as a result of the heating, aggravating the warping effect and reducing the image resolution.

As specific beamline time was pre-allocated, all screened blocks were mounted for beamline imaging. The blocks were then scored and sorted according to the number of artefacts found in the microCT screen. The sample screened with the least amount of visible artefacts (like cracks, bubbles or staining gradients) was sample “Durcupan-C” or “DC” and was therefore chosen for XNH imaging. A microCT image of sample DC can be seen in Figure 2a.

**Figure 1.**
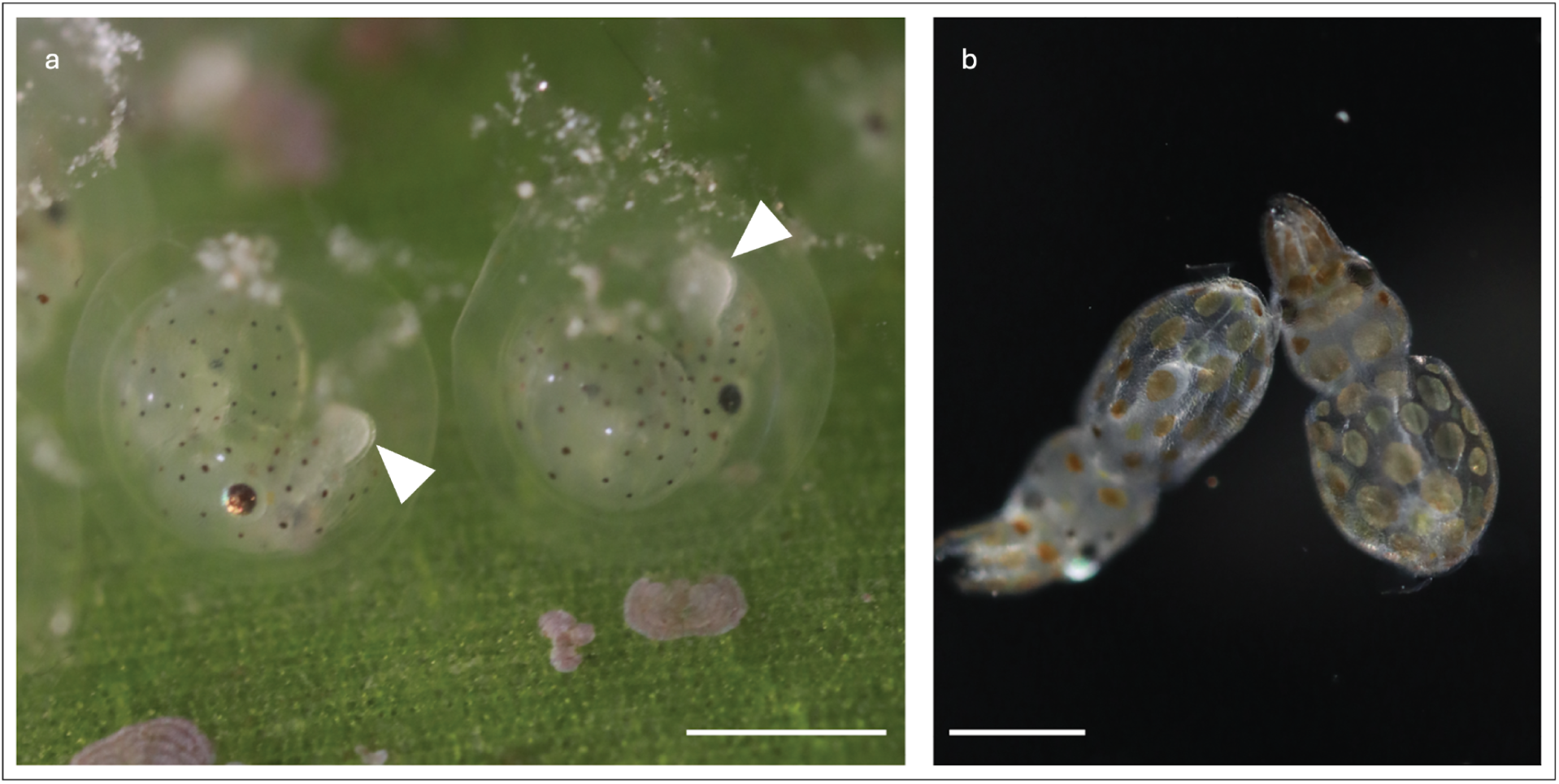
The Australian pygmy squid, Idiosepius hallami. - a) embryos possessing a small yolk sac (arrow head); b) hatchlings. scale bar: 1mm

**Figure 2.**
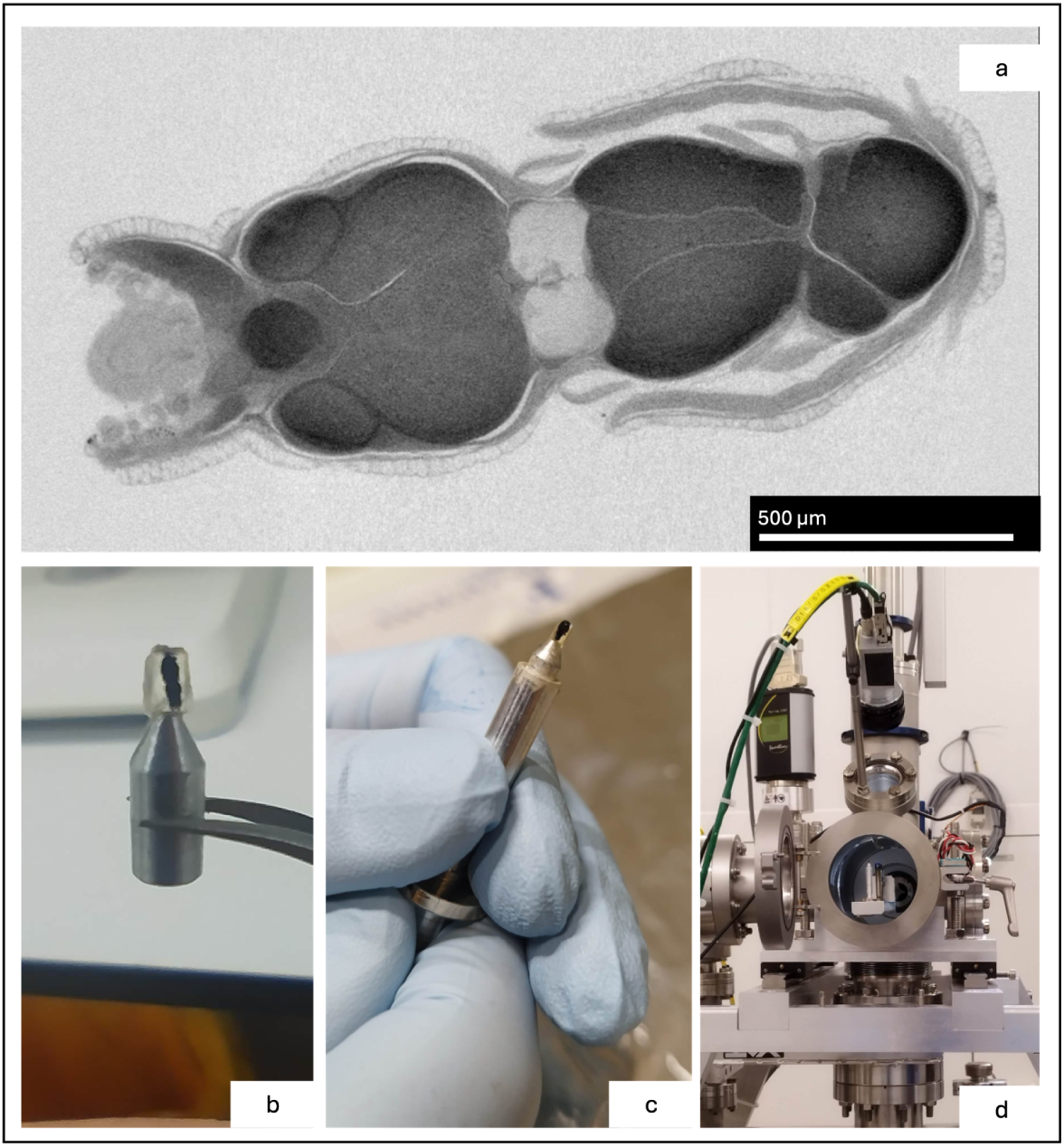
Sample preparation for X-ray nano holography. a) MicroCT of the Durcupan-C sample. b) Re-embedding of the Durcupan-C sample on the beamline-compatible aluminium stub. c) Fully prepared sample with smooth surrounding resin in the beamline holder; d) Sample loading into the vacuum chamber. (Figure included in the official ESRF experiment report) (Silva, 2024)

Samples were removed from the original resin block, trimmed to the minimal volume containing the animal and reattached on top of an aluminium stud (Figure 2b). A small drop of fresh resin was placed surrounding the trimmed specimen and polymerised upside down, creating a smooth bell-shaped coating that eliminates the sharp edges made by razor blade trimming that would lead to imaging artefacts. The aluminium stud was introduced in a sample holder (Figure 2c) and placed in the beamline chamber (Figure 2d) for the optimisation of the imaging parameters.

### Imaging parameters

Since neither Durcupan resin nor squid tissue had ever been imaged at the beamline ID16A, it was necessary to perform test scans at different resolutions and X-ray doses, checking the maximum achievable resolution that did not compromise resin stability and allowed the acquisition of the whole animal within the pre-allocated time. These test scans covered different areas, such as the arms and the brain (Figure 3). The scans were acquired aiming at a voxel size within the range of 40-150 nm. We concluded the tests by verifying that there was very little resin expansion or contraction, proving Durcupan to be a suitable resin for synchrotron imaging.

**Figure 3.**
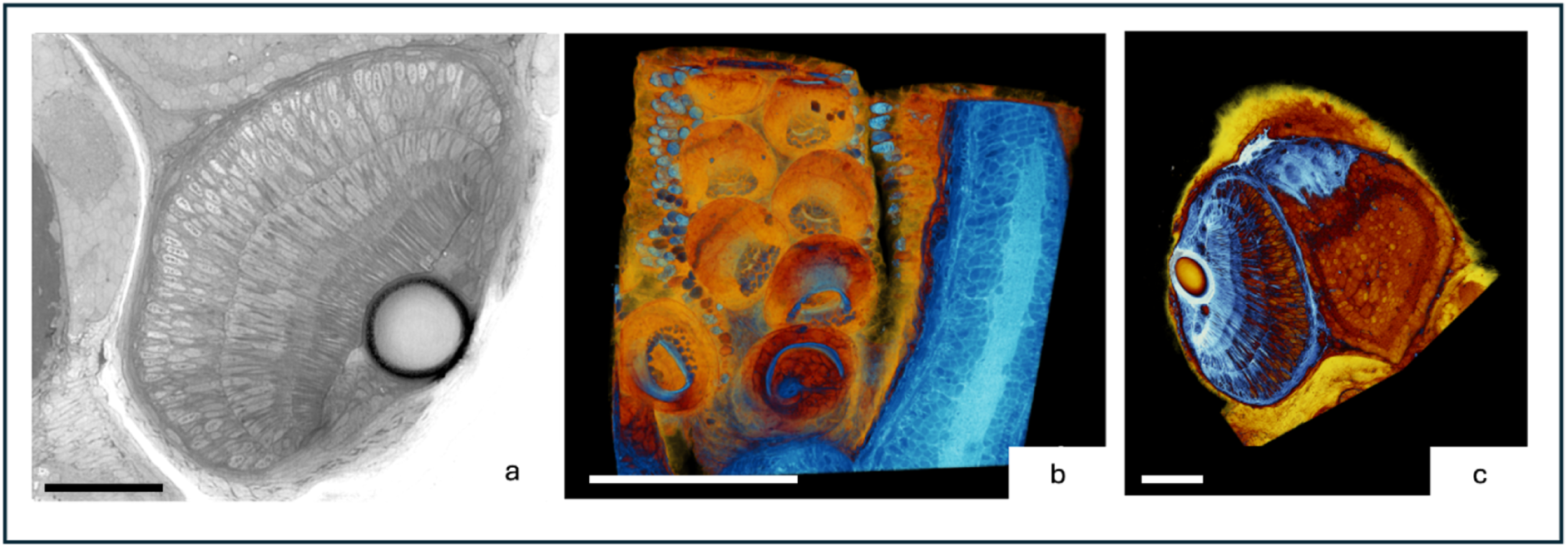
Test scans in the beamline ID16a. a) Test scan of the pygmy squid eye at a voxel size of 100nm. Approximate scale bar: 50 µm. b) and c) 3D rendering of test scans, b) a small portion of the pygmy squid arms, which show suckers and the underlying neural tracts, and c) a portion of the eye and the optic lobe (visual centre). The 3D rendering was acquired using the ParaView software and false colouring obtained with a custom LUT (Ahrens, Geveci and Law, 2005). Approximate scale bars: b) 100 µm and c) 50 µm. (Figure presented in the official ESRF experiment report) (Silva, 2024)

### Nano holotomography acquisition

The Durcupan-C sample was fully imaged at the nano-imaging beamline ID16A of the European Synchrotron - ESRF. The beamline endstation is located 185 meters from the source, providing a coherent nano-focused beam. This instrument is therefore optimised for coherent imaging, particularly for holographic tomography and X-ray fluorescence mapping. Kirkpatrick-Baez mirrors with a fixed curvature generate a spot size down to 13 nm at an energy of 33.6 keV (Julio Cesar da Silva et al., 2017) and a photon flux on the order of 10^11^ ph/s. The fixed pygmy squid sample was placed in the vacuum chamber of the endstation, which is fitted with a cryogenic rotation stage and an active stabilisation system, enabling precise positioning (Villar et al., 2018). After converting X-rays to visible light with an LSO-Tb scintillator, the images were recorded using a sCMOS sensor embedded in an Ximea camera.

The smallest voxel size that still allowed the imaging of the entire organism within the scheduled beamline time was chosen. To cover the whole pygmy squid, we acquired 40 tiled scans at eight vertical positions along the Z axis (with an overlap of 5-10%). Four to six round tiles overlapping by approximately 30% were selected and programmed for imaging at each of the eight levels. After reconstruction, each tile is a cylinder with a diameter and height of 300 µm and an isotropic voxel size of 125 nm (Figure 4a). We recorded four sets of 2,000 angular projections over 180° at different focus-to-sample-to-detector distances, enabling the acquisition of a full holo-tomographic scan. The exposure time for each projection was 0.28 seconds.

**Figure 4.**
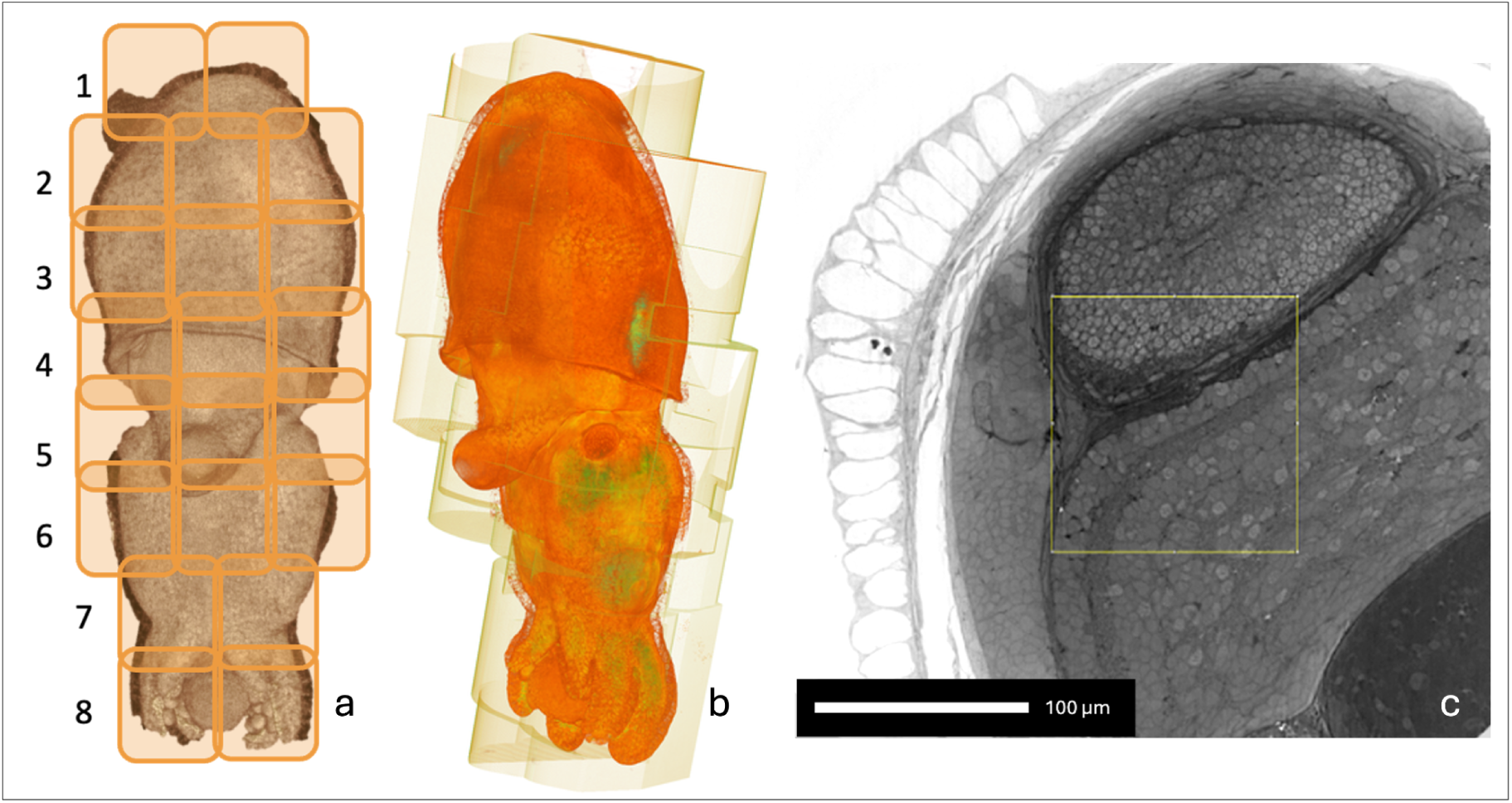
Scan set up and reconstruction. a) Virtual division of the pygmy squid in 8 levels, or scan tiles. b) rendering of reconstructed pygmy squid, showing the scan tile borders. False colouring with custom LUT using the Paraview software; c) scan after phase reconstruction. Approximate scale bar: 100 µm. (Silva, 2024)

### Image processing

Each projection was first processed for empty beam and optical distortion correction. Subsequently, the images acquired at different propagation distances for each angular projection were aligned and brought to the same magnification. The four aligned holograms were then combined to obtain a phase map through an iterative phase retrieval algorithm (Cloetens et al., 1999; Yu et al., 2018; Kuan et al., 2020). Since the pygmy squid was impregnated with osmium tetroxide, we used the value for the ratio between the refractive index decrement (δ) and the absorption index (β) corresponding to Os at 33.6 keV - 27 as a regularisation parameter during phase retrieval. The resulting phase maps were used to reconstruct the 3D volumes using filtered back projection. Given the increase in FOV with distance from the beam focus, we utilised this additional information to reconstruct volumes with extended FOV, which served as overlap regions for aligning and stitching the individual volumes (Figure 4b).

### Stitching

Each of the eight vertical levels was stitched using the software NRstitcher (Miettinen et al., 2019). Adjacent tile positions were initialised based on the stage positions recorded during scans in the metadata of the images. If a layer of adjacent tiles were well aligned, the adjacent vertical layer would be stitched to it afterwards. The range of grey values was set between -430 and 100. Due to the use of an extended FOV, the outer areas of a tile will have lower resolution, which can cause issues with stitching when the overlap occurs only in those areas. For this reason, after stitching the complete dataset (Figure 5a), additional scans were needed to cover gaps due to insufficient overlap.

**Figure 5.**
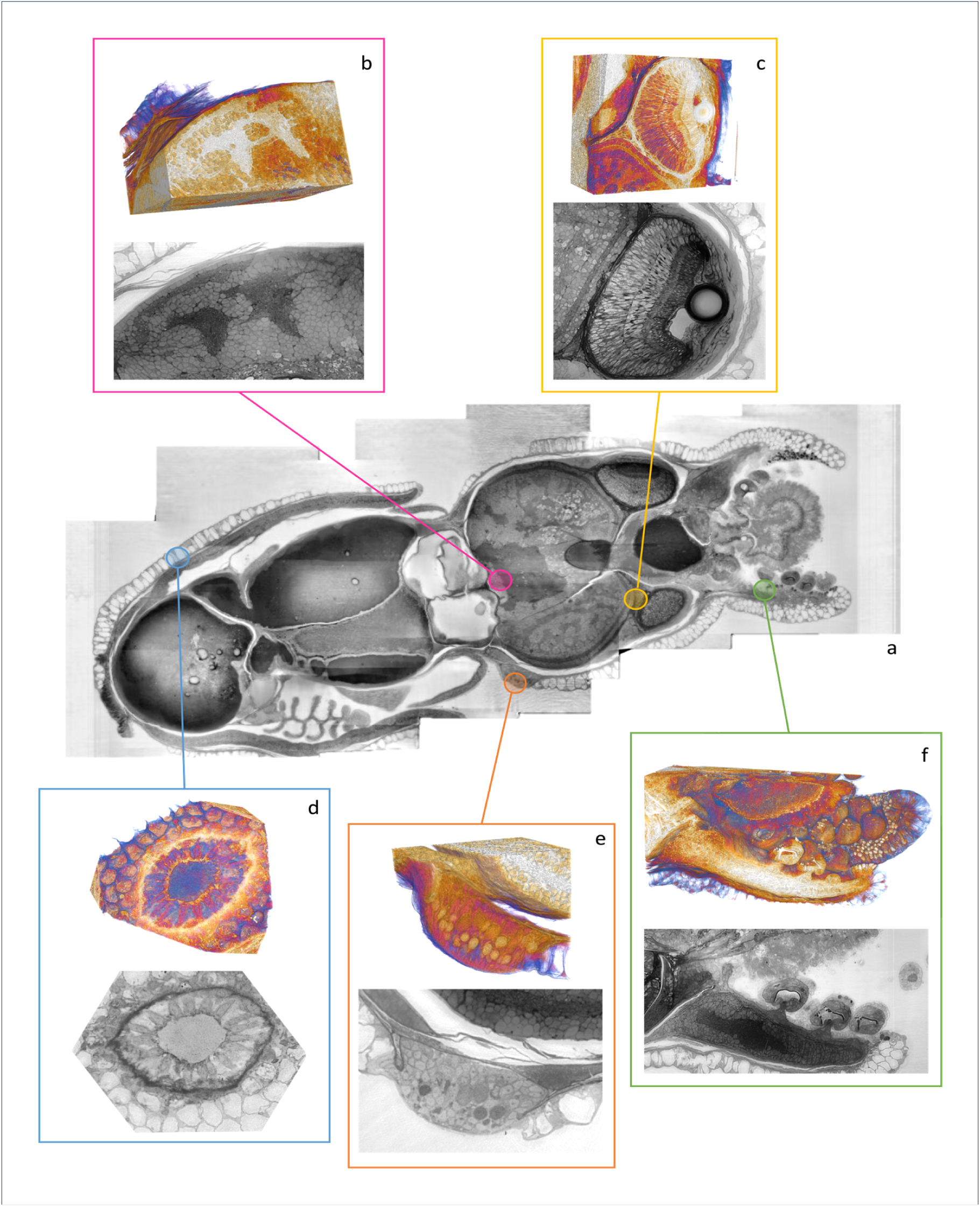
Overview of the dataset with 3D renderings. - a) 2D image of DC dataset. Not all regions highlighted in a-f are visible in this 2D image. b) 2D image and 3D rendering of vertical lobe complex; c) 2D image and 3D rendering of eye and part of an optic lobe; d) 2D image and 3D rendering of a chromatophore under the skin, showing radial muscles surrounding it; e) 2D image and 3D rendering of the olfactory organ of the pygmy squid; f) 2D image and 3D rendering of an arm, showing suckers attached to it. False colouring of 3D rendering obtained with a custom LUT in 3D slicer. (Silva, 2024)

### Atlas segmentation and identification of lobes, organs, and nerve tracts

The dataset was exported to N5 format, facilitating visualisation using CATMAID (Saalfeld et al., 2009a). The dataset level 3 (voxel size of 1 µm) was used to segment selected brain lobes using 3D slicer (Fedorov et al., 2012). The pygmy squid brain lobes were manually painted every 5 to 10 slices and then interpolated to create a continuous filling of the whole lobes. Interpolation was corrected in between painted slices when necessary. The lobes were identified according to previous cephalopod atlases (Yamamoto, Shimazaki and Shigeno, 2003; Koizumi et al., 2016; Chung, Kurniawan and Marshall, 2020; Montague et al., 2023), and the lobe boundaries were defined by the isolated areas of neuropil surrounded by somas, the somas being the outer boundary. Since the specimen is a pygmy squid hatchling, some lobes are not fully developed in the acquired dataset and could not be annotated. Each lobe was colour-labelled according to Table 2 to ease visualisation. Labels were exported to CATMAID, where they can be expanded to match full resolution.

**Table 2.** Cephalopod brain lobes and associated function. - Table adapted from Chung et al. (2020).

| Function | Complex | Lobe | Symmetry | Notes |
| --- | --- | --- | --- | --- |
| Learning and memory | Vertical lobe complex | Inferior frontal lobe |  |  |
|  |  | Superior frontal lobe |  |  |
|  |  | Posterior frontal lobe |  |  |
|  |  | Subvertical lobe |  |  |
|  |  | Vertical lobe |  |  |
| Higher motor control | Basal lobe complex | Anterior anterior basal lobe |  |  |
|  |  | Anterior Posterior basal lobe |  |  |
|  |  | Precommisural lobe |  |  |
|  |  | Dorsal basal lobe | Left and right |  |
|  |  | Interbasal lobe | Left and right |  |
|  |  | Median basal lobe |  |  |
|  |  | Lateral basal lobe | Left and right |  |
| Intermediate visual-motor center and olfaction | Optic tract complex | Peduncle lobe | Left and right |  |
|  |  | Olfactory lobe | Left and right |  |
|  |  | Dorsolateral lobe | Left and right |  |
| Vision | Optic lobe | Optic Lobe | Left and right |  |
| Arm and feeding control | Brachial lobe complex | Inferior buccal lobe |  |  |
|  |  | Superior buccal lobe |  |  |
|  |  | Brachial lobe |  | Was not separated from prebrachial lobe |
| Intermediate and lower motor center (locomotion control) | Pedal lobe complex | Anterior dorsal chromatophore lobe | Left and right | Difficult to tell apart in paralarvae |
|  |  | Anterior ventral chromatophore | Left and right | Difficult to tell apart in paralarvae |
|  |  | Anterior pedal lobe |  |  |
|  |  | Lateral pedal lobe | Left and right |  |
|  |  | Posterior pedal lobe |  |  |
| Intermediate motor center | Magnocellular lobe complex | Dorsal magnocellular lobe | Left and right |  |
|  |  | Ventral magnocellular lobe | Left and right |  |
|  |  | Posterior magnocellular lobe | Left and right |  |
| Lower motor center (locomotion) and mantle | Palliovisceral lobe complex | Palliovisceral lobe |  | No defined borders |
|  |  | Lateral ventral palliovisceral lobe | Left and right | No defined borders |
|  |  | Fin lobe | Left and right | No defined borders |
|  |  | Posterior chromatophore lobe | Left and right | No defined borders |
|  |  | Visceral lobe |  | No defined borders |

The open-source software CATMAID was used to visualise the dataset at full resolution (voxel size of 125 nm) and to trace nerve bundles using skeleton-based segmentation. Nerve tracts were labelled according to previous work (Shigeno and Yamamoto, 2002; Chung, Kurniawan and Marshall, 2020). 3D volumes of selected lobes were also created in CATMAID using alpha shape volumes to facilitate the location of the tracts.

Example subvolumes of the dataset were cropped in selected areas, such as the eye and optic lobe, vertical lobe, olfactory organ, chromatophore and arm. The individual renderings are displayed in Figure 5. These cropped volumes were visualised using the plugin Fiji (Schindelin et al., 2012), BigDataViewer (Pietzsch et al., 2015) and 3D rendered using 3D slicer (Fedorov et al., 2012).

Additionally, we implement an automated pipeline using the BiaPy library (Franco-Barranco et al., 2025) for (i) nuclei segmentation and (ii) nerve-tract segmentation and tracking across the full volume by adopting 3D Residual U-Net-based deep learning models. For both nuclei and nerve tracts, semantic probability maps are fused across tiles and connected components are obtained via morphological watershed. Outputs integrate into CATMAID for interactive proofreading and quality control.

## Results

### Overview of the squid XNH volume

The montaged volume (Figure 5a) encompasses the full body of the pygmy squid hatchling (dimensions 0.7 * 0.7 * 2 mm) at a voxel size of 125 nm, combining thirty-six initial 3D scans and four supplementary scans to address missing overlaps. While montaging was successful for most scans, a few artefacts were introduced by alignment and distortion errors at the interfaces. These artefacts do not interfere with the segmentation of large structures such as brain lobes, and nerve bundles remain traceable across adjacent scans. Ring artefacts, appearing as concentric rings producing stripes in the reconstructed synchrotron images, are a common issue (Vo, Atwood and Drakopoulos, 2018) that impacts mainly the dataset’s appearance.

According to the embryonic atlas of *Idiosepius* created by Yamamoto et al. (2003), the underdevelopment of certain brain lobes and the substantial yolk remaining in the digestive tract suggest that the specimen is timed between stages 27 and 29.

In particular, Yamamoto et al. (2003) noted that during stages 25 to 27, the arms neuropil (intrabrachial ganglia), the superior buccal lobe and the anterior subesophageal mass (comprising prebrachial lobe and brachial lobe) become anatomically distinguishable from the brain. By stage 27, the anterior subesophageal mass separates from the middle subesophageal mass (containing the anterior chromatophore lobe, anterior pedal lobe, posterior pedal lobe, and ventral magnocellular lobe). Figure 8 shows an evident separation of these two masses, supporting the classification of the specimen as being at least at stage 27. We can discern major brain lobes at this stage, although some are underdeveloped. The vertical lobe, for example, is very small compared to that of the adult (Figure 5b).

### Arms and tentacles

The arms are located in the most anterior region of the animal (Figure 5f). These highly innervated structures are essential for dexterous behaviours such as object manipulation, feeding and hunting. Within the arms, a well-demarcated axial neuropil is surrounded by somas and muscle bundles, which connect to suckers (Figure 5f). Additionally, very small tentacle primordia are visible, flanking the mouth of the animal (Figure 6). Squid tentacles are long and flexible appendages, mediating with the tentacular ballistic strike onto the fast-moving prey.

**Figure 6.**
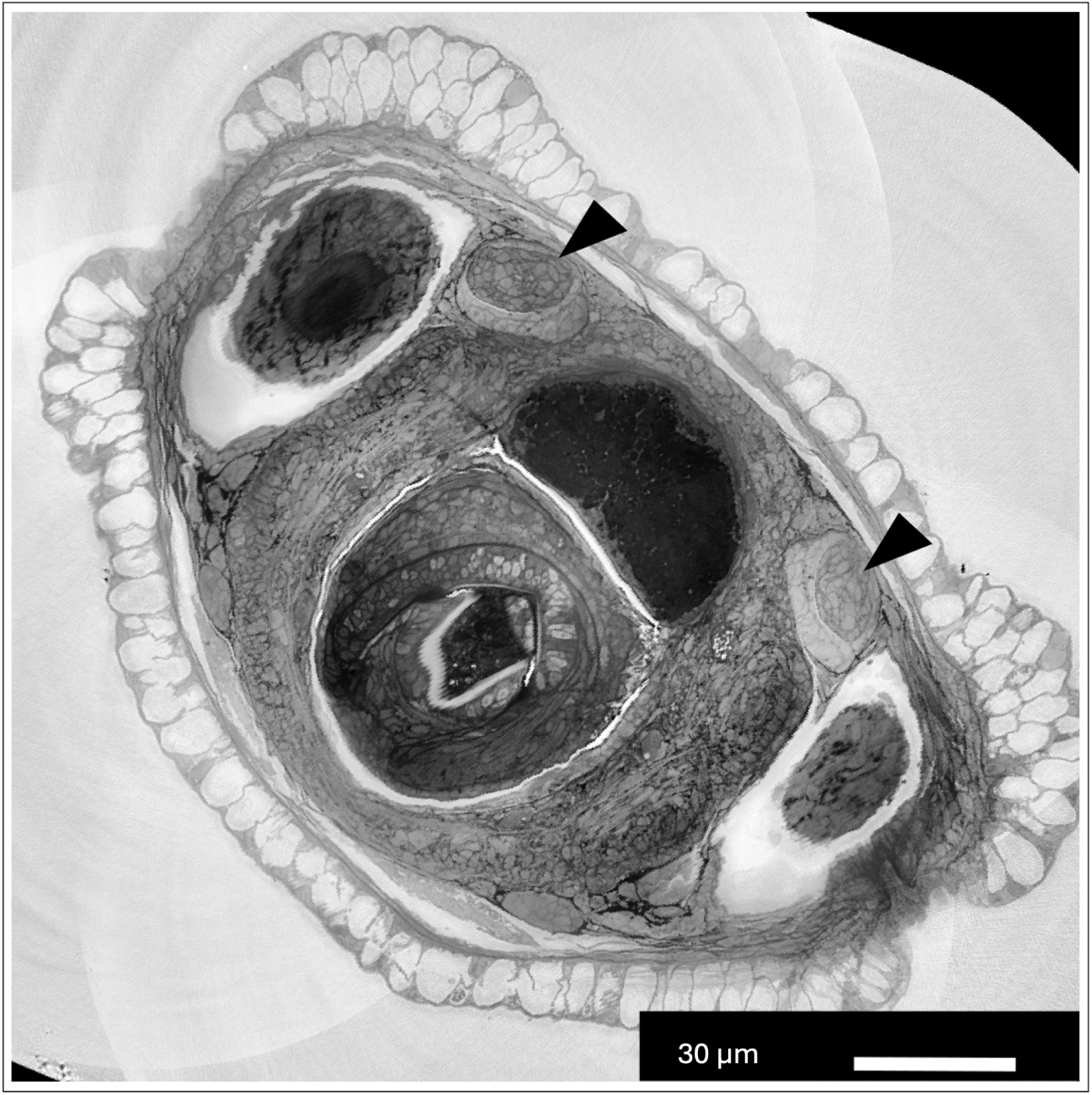
Tentacle primordia. - X-ray reconstructed photograph showing a virtual transversal section of the pygmy squid. Small tentacle primordia are visible (black arrows).

### Central brain and optic lobes

The head region of the pygmy squid contains the multi-lobed brain, the eyes and the buccal mass. The paired, camera-like eyes are situated superficially, adjacent to the optic lobes, and are connected to the brain via the optic nerves (Figure 5c). Their major constituents are clearly distinguishable: the iris, the spherical lens, the vitreous cavity, and the retina with its rhabdomeric photoreceptors. The optic lobes, which are responsible for image processing, constitute the largest part of the brain and were segmented alongside the lobes in the optic tract complex (peduncle, olfactory and dorsolateral lobes) (Figure 7e). The optic lobes are well-defined, bean-shaped lobes located on each side of the brain and are connected by optic tracts, which are also clearly distinguishable in the volume (Figure 7e and 7g).

**Figure 7.**
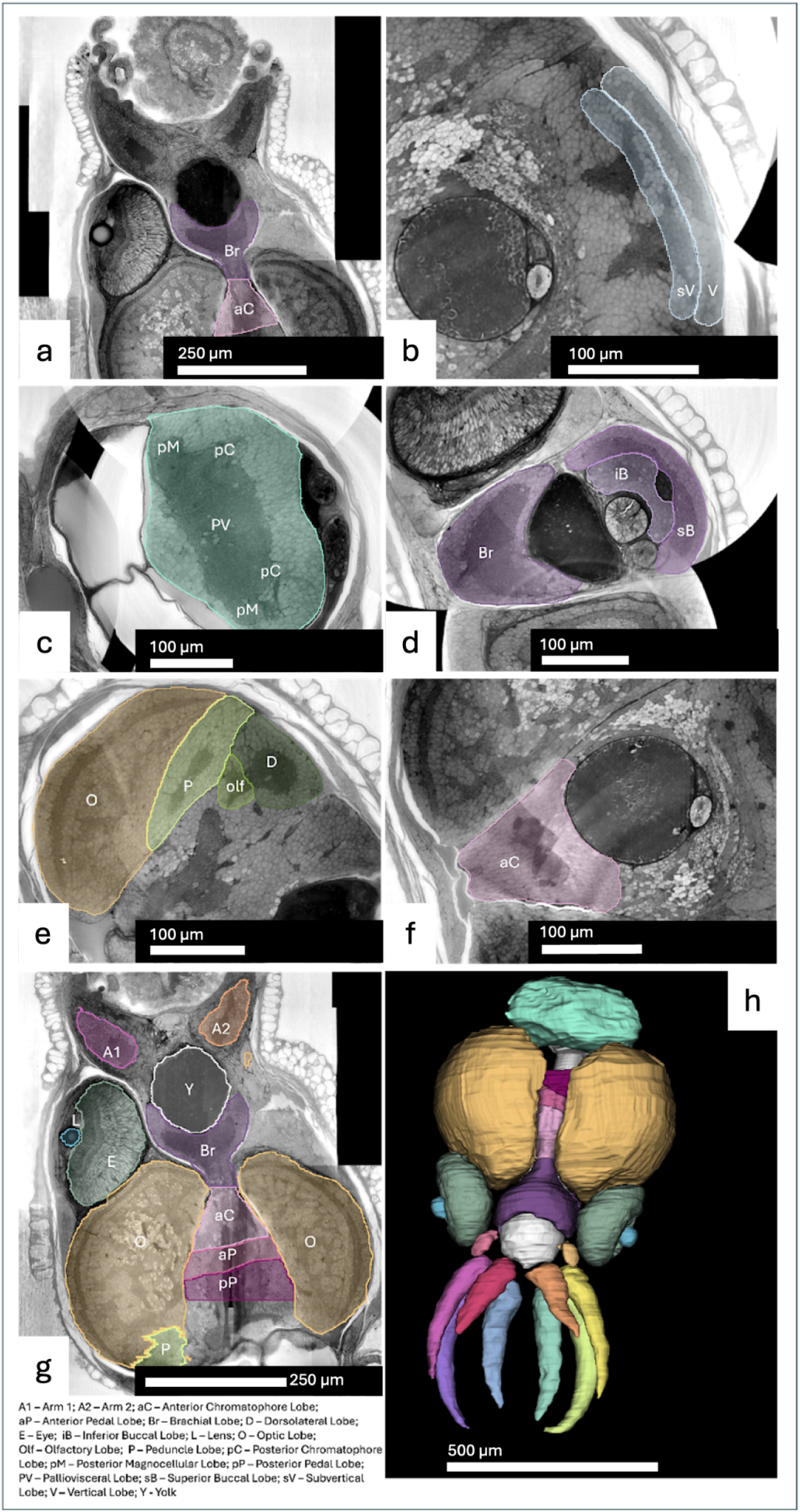
Overview of the dataset with segmented brain lobes. a-g) Reconstructed X-ray images of transverse sections of the pygmy squid brain and painted selected regions. a) anterior subesophageal and middle subesophageal masses, showing a clear separation between them; b) vertical and sub-vertical lobes; c) Palliovisceral lobe complex; d) Optic lobes and optic tract complexes comprising peduncle, olfactory and dorsolateral lobes; e) Brachial lobe complex, comprising brachial, inferior buccal and superior buccal lobes; f) Anterior chromatophore lobes. g) A few of the structures and brain lobes segmented; h) 3D rendering of structures and brain lobes segmented. (Preliminary data) (Silva, 2024)

The brachial lobe complex, responsible for controlling the arms and movements related to feeding, was segmented and separated into the brachial, inferior buccal and superior buccal lobes (Figure 7a).

The vertical lobe (VL), the associative learning centre of the pygmy squid, is located in the most dorsal part of the CNS (Figure 7b). Notably, unlike the dome-shaped VL of the adult (Chung and Marshall, 2017), the hatchling’s vertical lobe, where the neuropil forms a very thin plate surrounded by somas, remains. The vertical and sub-vertical lobes can be seen in Figure 7b.

The palliovisceral lobe complex is generally well-defined. However, individual lobes are difficult to distinguish at this resolution and developmental stage. Nonetheless, we estimated the regions corresponding to the posterior chromatophore, fin, visceral, lateral ventral palliovisceral and palliovisceral lobes (Figure 7c).

### Sensory organs

The olfactory organs (Figure 5e) are located on either side of the head, posterior and ventral to each eye, near the mantle edge. These chemosensory organs are composed of ciliated epithelial cells, mucus-producing cells, and various types of sensory cells (Emery, 1975; Mobley, Mahendra and Lucero, 2007; Polese, Bertapelle and Cosmo, 2016). The axons of these sensory neurons bundle into the olfactory nerve, which projects to the olfactory lobe (Figure 7e). To our knowledge, this is the first time this long projection from the olfactory organ to the olfactory lobe has been clearly visualised.

A pair of statocysts is situated beneath the brain, within which the statolith is clearly discernible. (Figure 8). These structures help maintain the balance and orientation of the pygmy squid by providing information on movement and acceleration. Additionally, they also detect infrasound (low-frequency vibrations)(Williamson and Budelmann, 1985; Hu et al., 2009).

**Figure 8.**
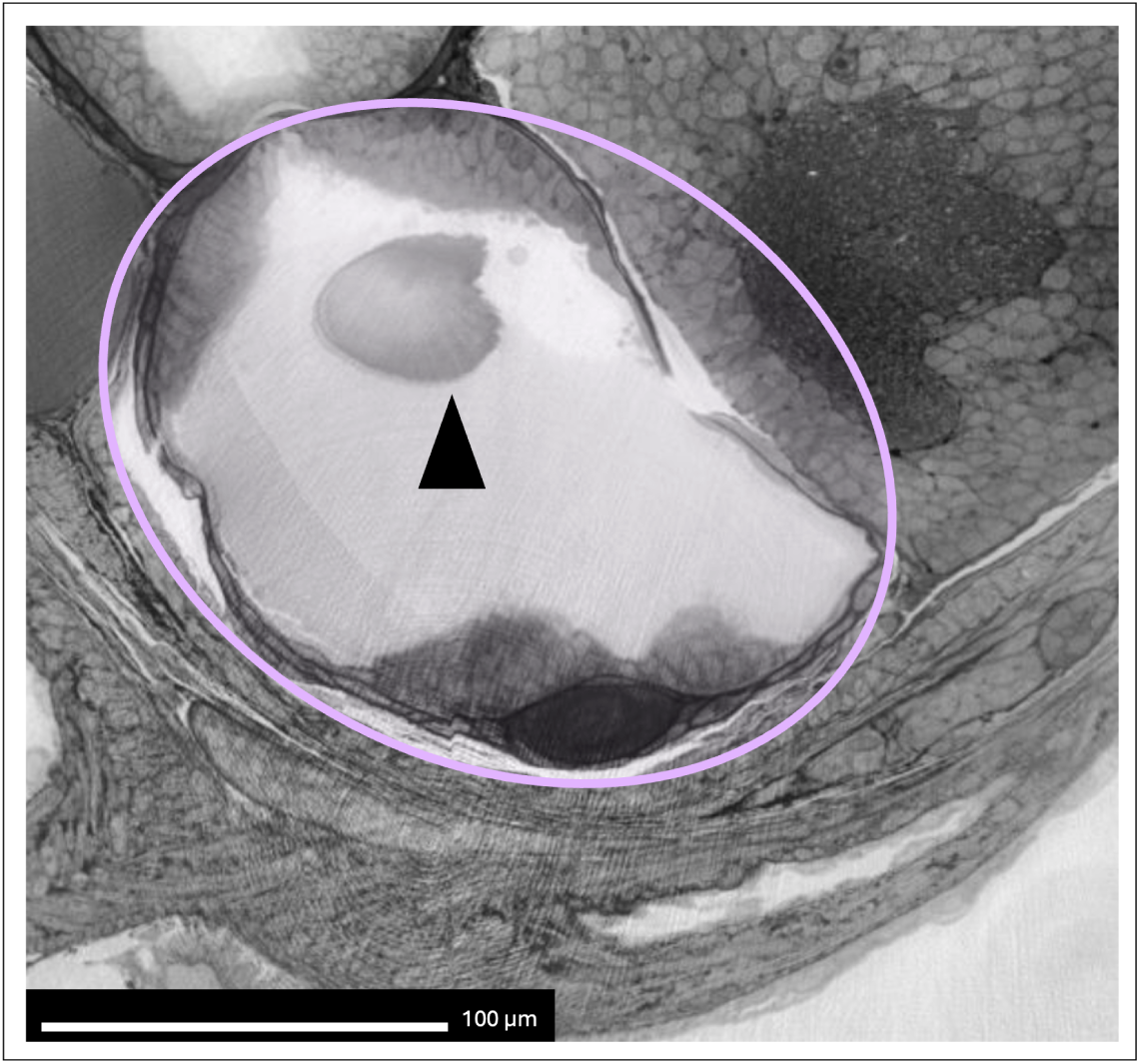
Statocysts and statolith. - Virtual slice of the pygmy squid volume transversal to the antero-posterior axis. One statocyst is surrounded by light purple, while the statolith is identified with a black arrow.

The mass right below the arrowhead is the “crista/cupula system”, which can detect the movement.

### Internal organs in the mantle

In the posterior region of the pygmy squid, a mantle encloses all internal organs. The stellate ganglia, which are connected to the palliovisceral lobe via the pallial nerves, are located on each side of the mantle (Figure 10). From the stellate ganglia, stellar nerves extend into the mantle, gills, branchial hearts, and digestive system are also visible in this dataset.

### Chromatophores

Chromatophores, located beneath the skin, are distributed across the mantle. At the highest magnification, and owing to the ability to the isotropic volumes in any arbitrary direction, the structure of individual chromatophores is revealed when slices are perpendicular to the skin. Each chromatophore consists of a large central sac containing the pigment, surrounded by muscle fibres that radiate outward and attach to the sac. These muscles contract or relax, controlling the precise degree of pigment dispersion, allowing the squid to match its surroundings during camouflage (Figure 10). Finally, the anterior chromatophore lobes in the pedal lobe complex were segmented (Figure 7f).

### Mapped neural projections

Three sets of nerve bundles were traced using CATMAID (Saalfeld et al., 2009b) as a proof of principle.

### Olfactory nerves

The first set innervates the olfactory organ, located near the mantle edge on each side of the squid, which is responsible for detecting water-borne odorants (Mobley, Mahendra and Lucero, 2007). Figure 9 illustrates the neural projection from one olfactory organ to the olfactory and dorsolateral lobes. Afferent axons from the olfactory receptor neurons bundle in the olfactory organ, enter the optic tract region, and project to the olfactory lobe, subsequently reaching the dorsolateral lobes (Messenger, 1979). These two small lobes are located in the optic tract complex, alongside the peduncle lobe, and form part of the olfactory system of the cephalopod. In this dataset, the olfactory nerve is clearly identified and can be traced from the periphery to the centre of the brain (Figure 9).

**Figure 9.**
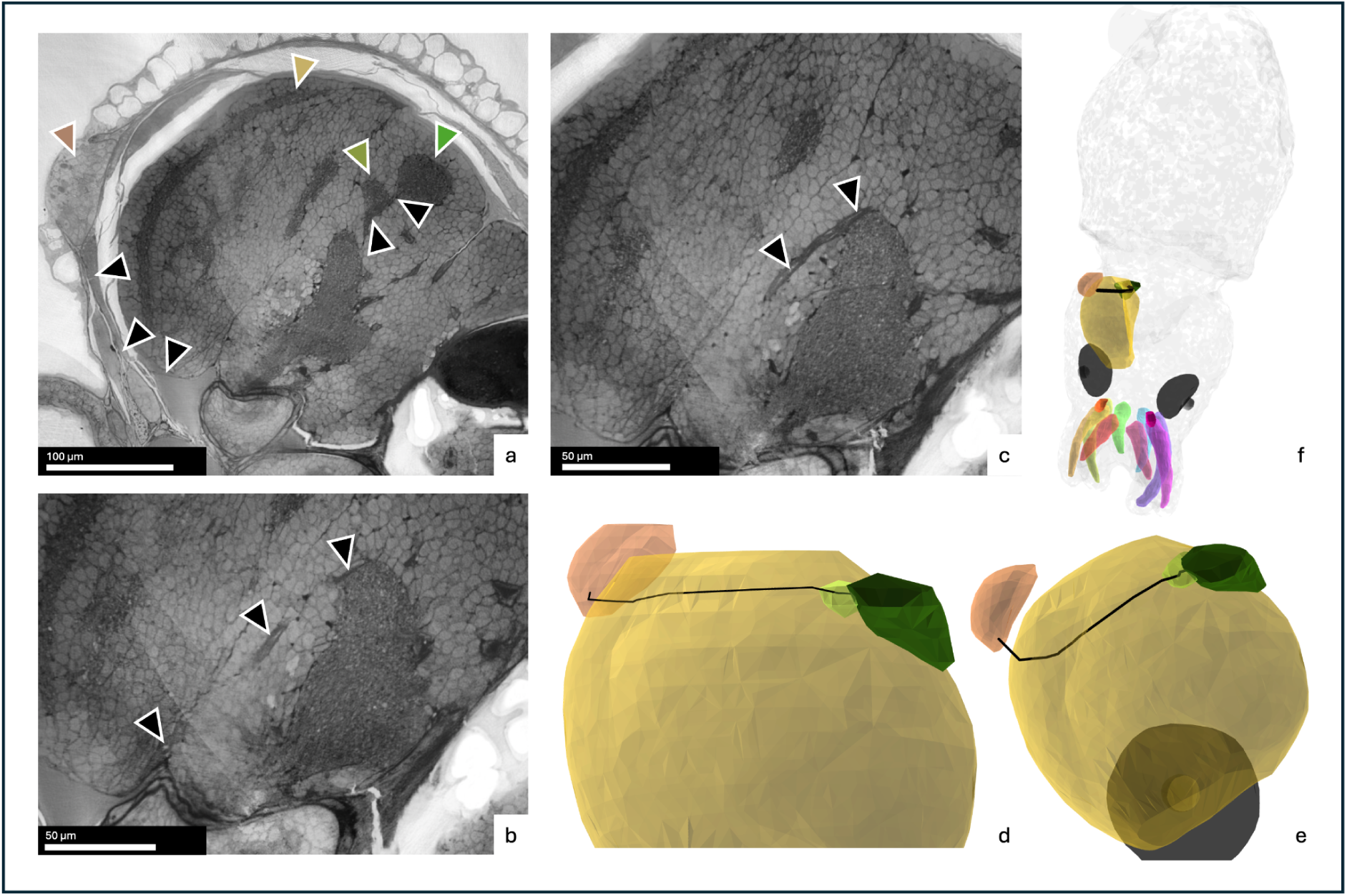
Projection from olfactory organ to olfactory lobe. - a-c) X-ray images of transverse sections of the pygmy squid brain. Brown arrow - olfactory organ; yellow arrow - optic lobe; light green arrow - olfactory lobe; dark green arrow - dorsolateral lobe; black arrows - olfactory nerve. The olfactory nerve enters the brain by running dorsally and anteriorly across the floor of the orbit (a-c). After reaching the optic tract complex, it enters the olfactory lobe and then the dorsolateral lobe (a). d-e) 3D renderings of olfactory organ, optic lobe, olfactory lobe, dorsolateral lobe, olfactory nerve and eye (in dark grey). Colours from renderings correspond to the same colours highlighted in arrows. f) rendering of whole pygmy squid, rendering illustrated in d and e, and colourful arms as a reference for orientation. Scale bars: a) 100 µm; b and c) 50 µm. (Silva, 2024)

### Innervation of the chromatophores

The second set of nerves corresponds to the neural projection of the chromatophore lobes to the chromatophores (Figure 10). Skin colour change and camouflage in cephalopods occur via the expansion and contraction of chromatophores. These specialised organs are spread throughout the cephalopod skin and are surrounded by 6-20 chromatophore muscles, which are innervated and receive input from the brain to expand the pigment sac inside the chromatophore, enabling rapid colour change within milliseconds (Cabej, 2012; Roger T Hanlon and Messenger, 2018). According to Chiao and Hanlon (2001), the camouflage pathway begins with visual input, which is processed through the eyes and the optic lobes, which in turn transmit signals to the lateral basal lobes, proceeding to the chromatophore lobes. These lobes will project efferent nerves to the skin where the chromatophores lie (Chiao and Hanlon, 2001). The anterior chromatophore lobe (pink arrow in Figure 10e) coordinates dynamic colour changes in the head and arms, while the posterior chromatophore lobe (located in the palliovisceral lobe complex, indicated with a blue arrow in Figure 10b) provides input to the mantle skin (Boycott, 1953; Messenger, 2001; Chung et al., 2023). As the XNH volume encompasses the entire animal, we could map efferent pathways from both the anterior and posterior chromatophore lobes.

**Figure 10.**
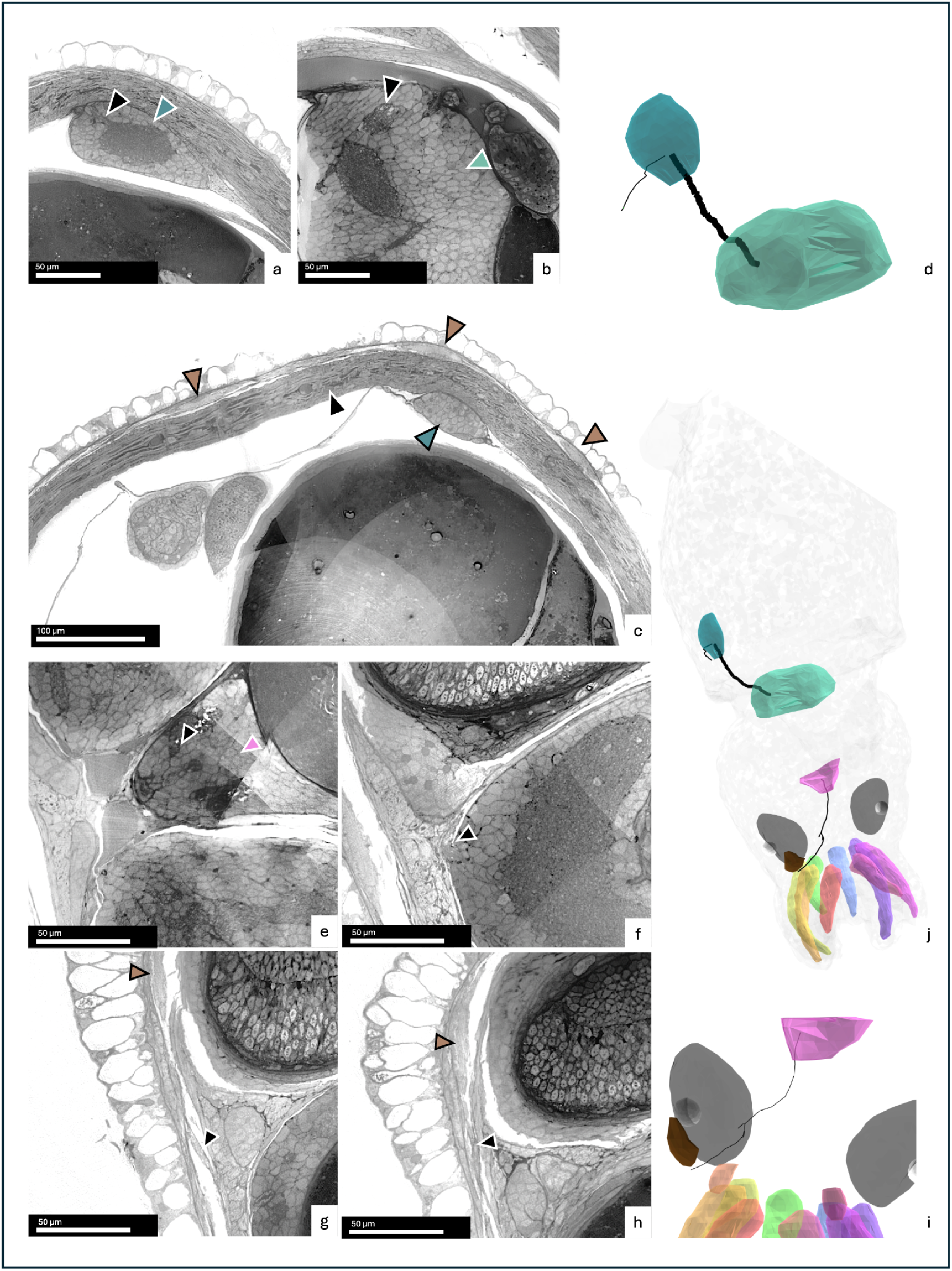
Projection from chromatophores to anterior and posterior chromatophore lobe. - a-c) X-ray images of transverse sections of the pygmy squid. Brown arrows - chromatophores; light blue arrow - palliovisceral lobe complex; dark blue arrow - stellate ganglion; black arrows - fibres from posterior chromatophore lobe to muscle underneath the cephalopod skin. Three chromatophores are near the area where the fibres are lost due to the dataset resolution. d) 3D rendering of the lobes and fibres described in a-c. e-h) X-ray images of transverse sections of the pygmy squid. Brown arrow - chromatophore; pink arrow - anterior chromatophore lobe; black arrows - fibres from anterior chromatophore lobe to muscle underneath the cephalopod skin, reaching close to the chromatophore. When the bundle disperses, the single fibres that reach the chromatophore lobe cannot be traced due to the dataset resolution. i) 3D rendering of the lobes and fibres described in e-i, with eyes (dark grey) and colourful arms for orientation reference. j) rendering of whole pygmy squid, rendering illustrated in d and i, and colourful arms as a reference for orientation. Scale bars: a-b) 50 µm; c) 100 µm; e-i) 50 µm. (Silva, 2024)

In the posterior chromatophore lobe (Figure 10), which is situated within the palliovisceral lobe complex, the efferent fibres travel through the pallial nerve to reach the stellate ganglion located outside the brain. From the stellate ganglion, these nerves reach the mantle skin, where the chromatophores are distributed.

Figure 10j illustrates the pathway of efferent fibres from the anterior chromatophore lobe that reach close to a chromatophore in the pygmy squid’s head.

### Innervation of the arms

The third set of nerves shows how the arms of the pygmy squid interconnect with one another and with the brain (Figure 11). Cephalopod arms are involved in a wide range of behaviours, including feeding and prey capture, locomotion, and reproduction (Vidal and Salvador, 2019). Squids possess eight arms and two tentacles, the latter of which are undeveloped at the hatchling stage. Each arm contains a central ganglion that is anatomically isolated from the brain. From the central arm neuropil, thick brachial nerves connect with the brachial lobe located in the anterior subesophageal mass (Figure 11c-d). The brachial nerves transmit information bidirectionally between the arms and the brain (Chang and Hale, 2023). From the brachial lobe, two thick nerves (known as the brachio-palliovisceral connectives) can be traced to the palliovisceral lobe complex (Figure 11a-b).

**Figure 11.**
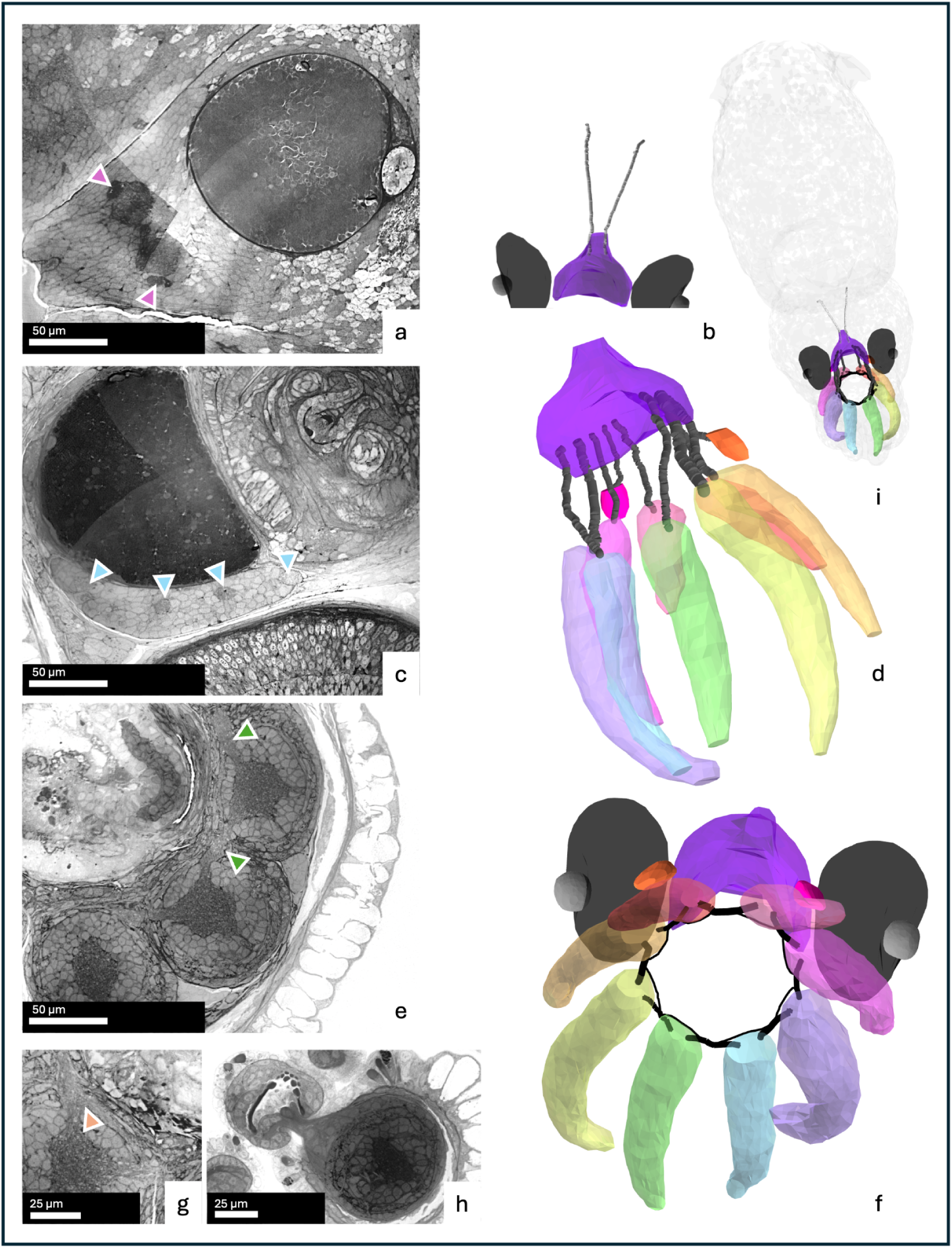
Connections between arms and brachial lobe, inter-arms and brachio-palliovisceral connective. - a, c, e, g and h) X-ray images of transverse sections of the pygmy squid. Pink arrows - brachio-palliovisceral connectives; light blue arrows - brachial nerves; green arrows - interbrachial commissures; orange arrow - fibres bypassing the arm, connecting to the adjacent interbrachial commissure and forming a second neural ring arrangement. a) Cross section of anterior chromatophore lobe showing the brachio-palliovisceral connective nerves; b) 3D rendering of the brachial lobe (dark purple) and brachio-palliovisceral connectives and eyes (dark grey) as a reference for orientation; c) brachial nerves in the brachial lobe; d) 3D rendering of the brachial lobe (dark purple) and brachial nerves connecting each arm and eyes (dark grey) as a reference for orientation; e and g) Cross section of arms showing central ganglia connected through interbrachial commissures (e) and interbrachial commissures connected through thinner nerves that bypass the arms (g); f) 3D rendering of the double ring configuration connecting the 8 arms and the brachial lobe (dark purple) and eyes (dark grey) as a reference for orientation; h) Sucker connected to arm; i) 3D rendering of whole pygmy squid, containing all the 3D renderings described above. Scale bars: a, c and e) 50 µm; g and h) 25 µm.(Silva, 2024)

At the base of each arm, the central ganglia project a lateral nerve bundle to each adjacent arm (Figure 11e-g). These nerve bundles, termed interbrachial commissures, are thick and short, directly linking one central ganglion to the next and forming a neural ring arrangement. A thinner nerve bundle branches from the centre of each interbrachial commissure (Figure 11g), bypassing the arm and connecting with the adjacent interbrachial commissure, forming a second, outer neural ring (Figure 11f). These pathways allow for interarm coordination (Chang and Hale, 2023). This configuration differs from the one described in *Octopus bimaculoides,* where each intramuscular nerve cord bypasses two adjacent arms (Kuuspalu, Cody and Hale, 2022). Additionally, small nerve bundles connecting the central arm neuropiles to the suckers are also evident (Figure 11h).

## Discussion

Cephalopods are formidable creatures, with one of the most sophisticated behavioural repertoires among invertebrates, making them an attractive model for neuroscience research (Reiter et al., 2018; Woo et al., 2023). While comparisons of convergent traits such as the visual system are well-established (Sweeney et al., 2007; Land and Nilsson, 2012; Nilsson, 2013; Napoli et al., 2022; Katz and Lyons, 2023), they often overlook the underlying neural architecture. The evolutionary divergence of cephalopods over 500 million years ago, combined with their remarkably analogous structures to vertebrates such as the camera eye and prehensile limbs, while sharing target pursuit strategies and limbed locomotion, offers an extraordinary opportunity to study convergent evolution in neural circuits in search of an optimal architectural principle common across phyla.

Although the pygmy squid *I. hallami* is small, this animal exhibits complex behaviours indicative of a fully functional nervous system (Kasugai, 2001; Nishiguchi et al., 2014; Koizumi et al., 2016; Sato et al., 2016; Reid and Strugnell, 2018). The optimised sample preparation protocol enabled whole-animal imaging using synchrotron XNH, yielding high contrast and minimal artefacts, while also being suitable for electron microscopy, supporting additional imaging of the same sample at higher resolution. Our protocol should be reproducible for other cephalopod species, with only minor modifications required to accommodate size differences.

Here, we introduced a whole-animal volume of an *Idiosepius halami* hatchling acquired by synchrotron X-ray imaging at 125 nm isotropic voxel size.

Before this experiment, Durcupan had never been tested in the synchrotron beamline ID16A. However, this is the resin of choice for the enhanced Focused Ion Beam Scanning Electron Microscopy technique (Xu et al., 2017), which we plan to implement in the same block. On the other hand, the epoxy resin Embed812 is widely available in the UK and was previously tested at the beamline (Kuan et al., 2020). For that reason, we tested both resins. The microCT screening showed prominent cracks in most samples embedded in Embed812 (Figure S1 a-e). Although some specimens embedded in Durcupan also presented cracks, these were less noticeable (Figure S1 f-j). Samples which had excessive external yolk were not chosen.

We segmented the brain lobes and traced a subset of key nerves connecting peripheral organs with the brain, such as the neural pathways linking the brain with the olfactory organs and the sparsely distributed chromatophores to the stellate ganglion and the brain. We further traced the neural and muscular innervation patterns of the arms, and indicated that the three-dimensional structure of the chromatophores can be analysed by isotropic slicing.

Advances in computer vision have now opened up the opportunity to generate a comprehensive projectome of the whole animal, facilitated by the automatic segmentation of nerve tracts and the nuclei that delineate the brain lobes. Although our primary focus was the nervous system, the volume of the whole pygmy squid hatchling offers further opportunities for researchers studying other aspects of cephalopod biology.

Despite the value of this dataset, there is room for improvement. First, future improvements to the stitching algorithm should more smoothly stitch each tile to each other, for in the present volume, distortion or misalignment artefacts can be observed between tiles. Furthermore, a few gaps persist between some of the acquired three-dimensional tiles, which can be addressed by reimaging the sample at those spots. Thankfully, further scans of small subvolumes can be integrated without altering the coordinates of the original scans, preserving original segmentations. Additionally, removal of ring artefacts will facilitate a more accurate segmentation, specifically for automated algorithms that aim to segment nuclei or nerve bundles in synchrotron datasets. Vo et al. (2018) propose several methods for ring artefact removal that can be implemented in the pygmy squid dataset. However, testing and optimising these methods will require significant effort.

In summary, we have generated a whole animal dataset that strikes a good balance between resolution and completeness, providing a robust foundation for future analyses. We segmented arms, organs, funnel, fins, and chromatophores, among other structures, alongside their nervous projections to/from the brain. Although individual neurons cannot be resolved at a voxel size of 125 nm, nerve bundles are clearly visible and traceable. Hence, we characterised the major afferents and efferents to the brain. The acquisition of the pygmy squid’s whole body volume represented a significant milestone for the synchrotron beamline ID16A, as it was, at that time, the largest biological volume imaged at a voxel size of 125 nm. By pushing the boundaries of current imaging technology, this work paved the way for the development of improved methods capable of acquiring larger, high-resolution datasets in a fraction of the time required by other techniques. Furthermore, the enthusiasm generated by this and similar volume acquisitions (Kuan et al., 2020) at beamline ID16A led to a proposal for a new beamline at the ESRF, dedicated entirely to connectomics.

## Data availability

The volume will be publicly accessible upon publication.

## Acknowledgements

We acknowledge the European Synchrotron (ESRF) for the provision of beamtime under proposal number LS-3187, and we would like to thank Dr Peter Cloetens for assistance and support in using beamline ID16A. We thank Amy Streets and Nicole Schieber at the Centre for Microscopy and Microanalysis (University of Queensland) for support in sample preparation. We thank Jayde Livingston for support in compiling the NR stitcher software used for stitching scans. We thank Ivana Henry for lab maintenance, and Christopher Barnes and Peter Hague for server maintenance.

Funded by a Wellcome Trust Investigator award to A. Cardona (Ref: 205038/Z/16/Z) and the MRC LMB core funding. We acknowledge funding support from the Australia and Pacific Science Foundation (APSF22032) to WSC, and from the European Commission through the ERC Brilliance grant (852455) to AP. We thank the staff of the Moreton Bay Research Station for logistical support. We also acknowledge the Quandamooka people as the Traditional Owners and their custodianship of the land on which Moreton Bay Research Station operates. We pay our respects to their ancestors and descendants, who continue cultural and spiritual connections to Country and recognise their valuable contributions to Australian and global society.

## References

Abbott, N.J., Williamson, R. and Maddock, L. (1995) “Cephalopod Neurobiology,” Oxford University Press, 75(3), pp. 767–767. Available at: 10.1017/s0025315400039217.

Ahrens, J., Geveci, B. and Law, C. (2005) “Visualization Handbook,” Part IX: Visualization Software and Frameworks, (IEEE Computer Graphics and Applications941989), pp. 717–731. Available at: 10.1016/b978-012387582-2/50038-1.

Albertin, C.B. and Katz, P.S. (2023) “Evolution of cephalopod nervous systems,” Current Biology, 33(20), pp. R1087–R1091. Available at: 10.1016/j.cub.2023.08.092.

Amodio, P., Boeckle, M., Schnell, A.K., Ostojíc, L., Fiorito, G. and Clayton, N.S. (2019) “Grow Smart and Die Young: Why Did Cephalopods Evolve Intelligence?,” Trends in Ecology & Evolution, 34(1), pp. 45–56. Available at: 10.1016/j.tree.2018.10.010.

Boycott, B.B. (1953) “THE CHROMATOPHORE SYSTEM OF CEPHALOPODS,” Proceedings of the Linnean Society of London, 164(2), pp. 235–240. Available at: 10.1111/j.1095-8312.1953.tb00688.x.

Boycott, B.B. (1961) “The functional organization of the brain of the cuttlefish Sepia officinalis,” Proceedings of the Royal Society of London. Series B. Biological Sciences, 153(953), pp. 503–534. Available at: 10.1098/rspb.1961.0015.

Boycott, B.B. (1965) “Learning in the Octopus,” Scientific American, 212(3), pp. 42–50. Available at: 10.1038/scientificamerican0365-42.

Boycott, B.B. and Young, J.Z. (1957) “Effects of interference with the vertical lobe on visual discriminations in Octopus vulgaris Lamarck,” Proceedings of the Royal Society of London. Series B - Biological Sciences, 146(925), pp. 439–459. Available at: 10.1098/rspb.1957.0023.

Byern, J. von, Rudoll, L., Cyran, N. and Klepal, W. (2008) “Histochemical characterization of the adhesive organ of three Idiosepius spp. species,” Biotechnic & Histochemistry, 83(1), pp. 29–46. Available at: 10.1080/10520290801999316.

Cabej, N.R. (2012) “Epigenetic Principles of Evolution,” Part II: Neural-Developmental Premises of Evolutionary Adaptation, pp. 327–365. Available at: 10.1016/b978-0-12-415831-3.00010-0.

Cajal, S.R. (1917) “Contribucion al conocimiento de la retina y centros opticos de los cefalopodos,” Trabajos del laboratorio de investigacious biologicas de la universidad de Madrid, (15), pp. 1–82. Available at: https://bipadi.ub.edu/digital/collection/atlesmed/id/103317.

Chang, W. and Hale, M.E. (2023) “Mechanosensory signal transmission in the arms and the nerve ring, an interarm connective, of Octopus bimaculoides,” iScience, 26(5), p. 106722. Available at: 10.1016/j.isci.2023.106722.

Chiao, C.-C. and Hanlon, R.T. (2001) “Cuttlefish camouflage: visual perception of size, contrast and number of white squares on artificial checkerboard substrata initiates disruptive coloration,” Journal of Experimental Biology, 204(12), pp. 2119–2125. Available at: 10.1242/jeb.204.12.2119.

Chung, W.-S., Kurniawan, N.D. and Marshall, N.J. (2020) “Toward an MRI-Based Mesoscale Connectome of the Squid Brain,” iScience, 23(1), p. 100816. Available at: 10.1016/j.isci.2019.100816.

Chung, W.-S., Kurniawan, N.D. and Marshall, N.J. (2022) “Comparative brain structure and visual processing in octopus from different habitats,” Current Biology, 32(1), pp. 97–110.e4. Available at: 10.1016/j.cub.2021.10.070.

Chung, W.-S., López-Galán, A., Kurniawan, N.D. and Marshall, N.J. (2023) “The brain structure and the neural network features of the diurnal cuttlefish Sepia plangon,” iScience, 26(1), p. 105846. Available at: 10.1016/j.isci.2022.105846.

Chung, W.-S. and Marshall, J. (2014) “Range-finding in squid using retinal deformation and image blur,” Current Biology, 24(2), pp. R64–R65. Available at: 10.1016/j.cub.2013.11.058.

Chung, W.-S. and Marshall, N.J. (2016) “Comparative visual ecology of cephalopods from different habitats,” Proceedings of the Royal Society B: Biological Sciences, 283(1838), p. 20161346. Available at: 10.1098/rspb.2016.1346.

Chung, W.-S. and Marshall, N.J. (2017) “Complex Visual Adaptations in Squid for Specific Tasks in Different Environments,” Frontiers in Physiology, 8, p. 105. Available at: 10.3389/fphys.2017.00105.

Cloetens, P., Ludwig, W., Baruchel, J., Dyck, D.V., Landuyt, J.V., Guigay, J.P. and Schlenker, M. (1999) “Holotomography: Quantitative phase tomography with micrometer resolution using hard synchrotron radiation x rays,” Applied Physics Letters, 75(19), pp. 2912–2914. Available at: 10.1063/1.125225.

Deryckere, A., Styfhals, R., Elagoz, A.M., Maes, G.E. and Seuntjens, E. (2021) “Identification of neural progenitor cells and their progeny reveals long distance migration in the developing octopus brain,” eLife, 10, p. e69161. Available at: 10.7554/elife.69161.

Emery, D.G. (1975) “The histology and fine structure of the olfactory organ of the squid Lolliguncula brevis blainville,” Tissue and Cell, 7(2), pp. 357–367. Available at: 10.1016/0040-8166(75)90011-7.

Fedorov, A., Beichel, R., Kalpathy-Cramer, J., Finet, J., Fillion-Robin, J.-C., Pujol, S., Bauer, C., Jennings, D., Fennessy, F., Sonka, M., Buatti, J., Aylward, S., Miller, J.V., Pieper, S. and Kikinis, R. (2012) “3D Slicer as an image computing platform for the Quantitative Imaging Network,” Magnetic Resonance Imaging, 30(9), pp. 1323–1341. Available at: 10.1016/j.mri.2012.05.001.

Fiorito, G., Planta, C. von and Scotto, P. (1990) “Problem solving ability of Octopus vulgaris lamarck (Mollusca, Cephalopoda),” Behavioral and Neural Biology, 53(2), pp. 217–230. Available at: 10.1016/0163-1047(90)90441-8.

Fiorito, G. and Scotto, P. (1992) “Observational Learning in Octopus vulgaris,” Science, 256(5056), pp. 545–547. Available at: 10.1126/science.256.5056.545.

Franco-Barranco, D., Román, J.A.A.-S., Hidalgo-Cenalmor, I., Backová, L., González-Marfil, A., Caporal, C., Chessel, A., Gómez-Gálvez, P., Escudero, L.M., Wei, D., Muñoz-Barrutia, A. and Arganda-Carreras, I. (2025) “BiaPy: accessible deep learning on bioimages,” Nature Methods, 22(6), pp. 1124–1126. Available at: 10.1038/s41592-025-02699-y.

Genoud, C., Titze, B., Graff-Meyer, A. and Friedrich, R.W. (2018) “Fast Homogeneous En Bloc Staining of Large Tissue Samples for Volume Electron Microscopy,” Frontiers in Neuroanatomy, 12, p. 76. Available at: 10.3389/fnana.2018.00076.

Hanlon, R. (2007) “Cephalopod dynamic camouflage.,” Current biology : CB, 17(11), pp. R400–4. Available at: 10.1016/j.cub.2007.03.034.

Hanlon, Roger T. and Messenger, J.B. (2018) Cephalopod Behaviour. Cambridge: Cambridge University Press (Cambridge Core). Available at: 10.1017/9780511843600.

Hanlon, Roger T and Messenger, J.B. (2018) “Cephalopod Behaviour,” pp. 45–73. Available at: 10.1017/9780511843600.005.

Hochner, B. and Glanzman, D.L. (2016) “Evolution of highly diverse forms of behavior in molluscs,” Current Biology, 26(20), pp. R965–R971. Available at: 10.1016/j.cub.2016.08.047.

How, M.J., Norman, M.D., Finn, J., Chung, W.-S. and Marshall, N.J. (2017) “Dynamic Skin Patterns in Cephalopods,” Frontiers in Physiology, 8, p. 393. Available at: 10.3389/fphys.2017.00393.

Hu, M.Y., Yan, H.Y., Chung, W.-S., Shiao, J.-C. and Hwang, P.-P. (2009) “Acoustically evoked potentials in two cephalopods inferred using the auditory brainstem response (ABR) approach,” Comparative Biochemistry and Physiology Part A: Molecular & Integrative Physiology, 153(3), pp. 278–283. Available at: 10.1016/j.cbpa.2009.02.040.

Huffard, C.L., Caldwell, R.L. and Boneka, F. (2008) “Mating behavior of Abdopus aculeatus (d’Orbigny 1834) (Cephalopoda: Octopodidae) in the wild,” Marine Biology, 154(2), pp. 353–362. Available at: 10.1007/s00227-008-0930-2.

Jereb, P. and Roper, C.F.E. (2005) Cephalopods of the world. An annotated and illustrated catalogue of cephalopod species known to date. Edited by F. and A.E. and P. Division. Rome: FAO (FAO Species Catalogue for Fishery Purposes, 1020-8682-No. 04 Vol.1). Available at: https://openknowledge.fao.org/handle/20.500.14283/a0150e.

Jereb, P. and Roper, C.F.E. (2010) Cephalopods of the world. An annotated and illustrated catalogue of cephalopod species known to date. Edited by F. and A.M. Division. Roma: Food and Agriculture Organization of the United Nations (FAO Species Catalogue for Fishery Purposes, 1020-8682-No. 04 Vol.2). Available at: https://openknowledge.fao.org/handle/20.500.14283/i1920e.

Jereb, P., Roper, C.F.E., Norman, M. and Finn, J.K. (2016) Cephalopods of the World. An Annotated and Illustrated Catalogue of Cephalopod Species Known to Date. Edited by F. species catalogue for fishery purposes. Roma: Food and Agriculture Organization of the United Nations. Available at: https://openknowledge.fao.org/handle/20.500.14283/i3489e.

Jozet-Alves, C., Schnell, A.K. and Clayton, N.S. (2023) “Cephalopod learning and memory,” Current Biology, 33(20), pp. R1091–R1095. Available at: 10.1016/j.cub.2023.08.013.

Kasugai, T. (2001) “Feeding behaviour of the Japanese pygmy cuttlefish Idiosepius paradoxus (Cephalopoda: Idiosepiidae) in captivity: evidence for external digestion?,” Journal of the Marine Biological Association of the United Kingdom, 81(6), pp. 979–981. Available at: 10.1017/s0025315401004933.

Kasugai, T. and Segawa, S. (2005) “Life cycle of the Japanese Pygmy Squid Idiosepius Paradoxus (Cephalopoda: Idiosepiidae) in the Zostera Beds of the Temperate Coast of Central Honshu, Japan.”

Katz, P.S. and Lyons, D.C. (2023) “Cephalopod vision: How to build a better eye,” Current Biology, 33(1), pp. R27–R30. Available at: 10.1016/j.cub.2022.11.054.

Koizumi, M., Shigeno, S., Mizunami, M. and Tanaka, N.K. (2016) “Three-dimensional brain atlas of pygmy squid, Idiosepius paradoxus, revealing the largest relative vertical lobe system volume among the cephalopods,” Journal of Comparative Neurology, 524(10), pp. 2142–2157. Available at: 10.1002/cne.23939.

Kuan, A.T., Phelps, J.S., Thomas, L.A., Nguyen, T.M., Han, J., Chen, C.-L., Azevedo, A.W., Tuthill, J.C., Funke, J., Cloetens, P., Pacureanu, A. and Lee, W.-C.A. (2020) “Dense neuronal reconstruction through X-ray holographic nano-tomography,” Nature Neuroscience, 23(12), pp. 1637–1643. Available at: 10.1038/s41593-020-0704-9.

Kuuspalu, A., Cody, S. and Hale, M.E. (2022) “Multiple nerve cords connect the arms of octopuses, providing alternative paths for inter-arm signaling,” Current Biology, 32(24), pp. 5415–5421.e3. Available at: 10.1016/j.cub.2022.11.007.

Land, M.F. and Nilsson, D.-E. (2012) “Animal Eyes,” pp. 72–93. Available at: 10.1093/acprof:oso/9780199581139.003.0004.

Lin, C.-Y., Tsai, Y.-C. and Chiao, C.-C. (2017) “Quantitative Analysis of Dynamic Body Patterning Reveals the Grammar of Visual Signals during the Reproductive Behavior of the Oval Squid Sepioteuthis lessoniana,” Frontiers in Ecology and Evolution, 5, p. 30. Available at: 10.3389/fevo.2017.00030.

Lu, X., Wu, Y., Schalek, R.L., Meirovitch, Y., Berger, D.R. and Lichtman, J.W. (2023) “A Scalable Staining Strategy for Whole-Brain Connectomics,” bioRxiv, p. 2023.09.26.558265. Available at: 10.1101/2023.09.26.558265.

Lu, Z., Xu, C.S., Hayworth, K.J., Pang, S., Shinomiya, K., Plaza, S.M., Scheffer, L.K., Rubin, G.M., Hess, H.F., Rivlin, P.K. and Meinertzhagen, I.A. (2022) “En bloc preparation of Drosophila brains enables high-throughput FIB-SEM connectomics,” Frontiers in Neural Circuits, 16, p. 917251. Available at: 10.3389/fncir.2022.917251.

Marini, G., Sio, F.D., Ponte, G. and Fiorito, G. (2017) “Learning and Memory: A Comprehensive Reference,” pp. 441–462. Available at: 10.1016/b978-0-12-809324-5.21024-9.

Marshall, N.J. and Messenger, J.B. (1996) “Colour-blind camouflage,” Nature, 382(6590), pp. 408–409. Available at: 10.1038/382408b0.

Messenger, J.B. (1979) “The nervous system of Loligo IV. The peduncle and olfactory lobes,” Philosophical Transactions of the Royal Society of London. B, Biological Sciences, 285(1008), pp. 275–309. Available at: 10.1098/rstb.1979.0007.

Messenger, J.B. (2001) “Cephalopod chromatophores: neurobiology and natural history,” Biological Reviews, 76(4), pp. 473–528. Available at: 10.1017/s1464793101005772.

Miettinen, A., Oikonomidis, I.V., Bonnin, A. and Stampanoni, M. (2019) “NRStitcher: non-rigid stitching of terapixel-scale volumetric images,” Bioinformatics, 35(24), pp. 5290–5297. Available at: 10.1093/bioinformatics/btz423.

Mikula, S. and Denk, W. (2015) “High-resolution whole-brain staining for electron microscopic circuit reconstruction,” Nature Methods, 12(6), pp. 541–546. Available at: 10.1038/nmeth.3361.

Mobley, A.S., Mahendra, G. and Lucero, M.T. (2007) “Evidence for multiple signaling pathways in single squid olfactory receptor neurons,” Journal of Comparative Neurology, 501(2), pp. 231–242. Available at: 10.1002/cne.21230.

Montague, T.G., Rieth, I.J., Gjerswold-Selleck, S., Garcia-Rosales, D., Aneja, S., Elkis, D., Zhu, N., Kentis, S., Rubino, F.A., Nemes, A., Wang, K., Hammond, L.A., Emiliano, R., Ober, R.A., Guo, J. and Axel, R. (2023) “A brain atlas for the camouflaging dwarf cuttlefish, Sepia bandensis,” Current Biology, 33(13), pp. 2794–2801.e3. Available at: 10.1016/j.cub.2023.06.007.

Napoli, F.R., Daly, C.M., Neal, S., McCulloch, K.J., Zaloga, A.R., Liu, A. and Koenig, K.M. (2022) “Cephalopod retinal development shows vertebrate-like mechanisms of neurogenesis,” Current Biology, 32(23), pp. 5045–5056.e3. Available at: 10.1016/j.cub.2022.10.027.

Nilsson, D.-E. (2013) “Eye evolution and its functional basis,” Visual Neuroscience, 30(1–2), pp. 5–20. Available at: 10.1017/s0952523813000035.

Nishiguchi, M.K., Nabhitabhata, J., Moltschaniwskyj, N.A. and Boletzky, S.V. (2014) “A review of the pygmy squid idiosepius: Perspectives emerging from an ‘inconspicuous’ Cephalopod,” Vie et Milieu, 64(December), pp. 23--34.

Nixon, M. and Young, J.Z. (2003) The Brains and Lives of Cephalopods. Edited by O.U. Press. Oxford, UK.

Osorio, D., Ménager, F., Tyler, C.W. and Darmaillacq, A.-S. (2022) “Multi-level control of adaptive camouflage by European cuttlefish,” Current Biology, 32(11), pp. 2556–2562.e2. Available at: 10.1016/j.cub.2022.04.030.

Pietzsch, T., Saalfeld, S., Preibisch, S. and Tomancak, P. (2015) “BigDataViewer: visualization and processing for large image data sets,” Nature Methods, 12(6), pp. 481–483. Available at: 10.1038/nmeth.3392.

Polese, G., Bertapelle, C. and Cosmo, A.D. (2016) “Olfactory organ of Octopus vulgaris: morphology, plasticity, turnover and sensory characterization,” Biology Open, 5(5), pp. 611–619. Available at: 10.1242/bio.017764.

Ponte, G., Taite, M., Borrelli, L., Tarallo, A., Allcock, A.L. and Fiorito, G. (2021) “Cerebrotypes in Cephalopods: Brain Diversity and Its Correlation With Species Habits, Life History, and Physiological Adaptations,” Frontiers in Neuroanatomy, 14, p. 565109. Available at: 10.3389/fnana.2020.565109.

Pungor, J.R., Allen, V.A., Songco-Casey, J.O. and Niell, C.M. (2023) “Functional organization of visual responses in the octopus optic lobe,” Current Biology, 33(13), pp. 2784–2793.e3. Available at: 10.1016/j.cub.2023.05.069.

Pungor, J.R. and Niell, C.M. (2023) “The neural basis of visual processing and behavior in cephalopods,” Current Biology, 33(20), pp. R1106–R1118. Available at: 10.1016/j.cub.2023.08.093.

Reid, A.L. and Strugnell, J.M. (2018) “A new pygmy squid, Idiosepius hallami n. sp. (Cephalopoda: Idiosepiidae) from eastern Australia and elevation of the southern endemic ‘notoides’ clade to a new genus, Xipholeptos n. gen.,” Zootaxa, 4369(4), pp. 451–486. Available at: 10.11646/zootaxa.4369.4.1.

Reiter, S., Hülsdunk, P., Woo, T., Lauterbach, M.A., Eberle, J.S., Akay, L.A., Longo, A., Meier-Credo, J., Kretschmer, F., Langer, J.D., Kaschube, M. and Laurent, G. (2018) “Elucidating the control and development of skin patterning in cuttlefish,” Nature, 562(7727), pp. 361–366. Available at: 10.1038/s41586-018-0591-3.

Saalfeld, S., Cardona, A., Hartenstein, V. and Tomancak, P. (2009a) “CATMAID: Collaborative annotation toolkit for massive amounts of image data,” Bioinformatics, 25(15), pp. 1984--1986. Available at: 10.1093/bioinformatics/btp266.

Sampaio, E., Sridhar, V.H., Francisco, F.A., Nagy, M., Sacchi, A., Strandburg-Peshkin, A., Nührenberg, P., Rosa, R., Couzin, I.D. and Gingins, S. (2024) “Multidimensional social influence drives leadership and composition-dependent success in octopus–fish hunting groups,” Nature Ecology & Evolution, 8(11), pp. 2072–2084. Available at: 10.1038/s41559-024-02525-2.

Sato, N., Takeshita, F., Fujiwara, E. and Kasugai, T. (2016) “Japanese pygmy squid (Idiosepius paradoxus) use ink for predation as well as for defence,” Marine Biology, 163(3), p. 56. Available at: 10.1007/s00227-016-2833-y.

Schindelin, J., Arganda-Carreras, I., Frise, E., Kaynig, V., Longair, M., Pietzsch, T., Preibisch, S., Rueden, C., Saalfeld, S., Schmid, B., Tinevez, J.-Y., White, D.J., Hartenstein, V., Eliceiri, K., Tomancak, P. and Cardona, A. (2012) “Fiji: an open-source platform for biological-image analysis,” Nature Methods, 9(7), pp. 676–682. Available at: 10.1038/nmeth.2019.

Schnell, A.K., Amodio, P., Boeckle, M. and Clayton, N.S. (2021) “How intelligent is a cephalopod? Lessons from comparative cognition,” Biological Reviews, 96(1), pp. 162–178. Available at: 10.1111/brv.12651.

Schnell, A.K. and Clayton, N.S. (2019) “Cephalopod cognition,” Current Biology, 29(15), pp. R726–R732. Available at: 10.1016/j.cub.2019.06.049.

Shigeno, S., Andrews, P.L.R., Ponte, G. and Fiorito, G. (2018) “Cephalopod Brains: An Overview of Current Knowledge to Facilitate Comparison With Vertebrates,” Frontiers in Physiology, 9, p. 952. Available at: 10.3389/fphys.2018.00952.

Shigeno, S. and Yamamoto, M. (2002) “Organization of the nervous system in the pygmy cuttlefish, Idiosepius paradoxus ortmann (Idiosepiidae, cephalopoda),” Journal of Morphology, 254(1), pp. 65–80. Available at: 10.1002/jmor.10020.

Silva, A.C.J.C.D. (2024) Exploring the Cephalopod Brain: Towards Mapping the First Whole Brain Connectome and Whole Animal Projectome of a Pygmy Squid. Apollo - University of Cambridge Repository.

Silva, J.C. da, Pacureanu, A., Yang, Y., Fus, F., Hubert, M., Bloch, L., Salome, M., Bohic, S. and Cloetens, P. (2017) “High-energy cryo x-ray nano-imaging at the ID16A beamline of ESRF,” X-Ray Nanoimaging: Instruments and Methods III, 10389, p. 103890F. Available at: 10.1117/12.2275739.

Song, K., Feng, Z. and Helmstaedter, M. (2023) “High-contrast en bloc staining of mouse whole-brain and human brain samples for EM-based connectomics,” Nature Methods, 20(6), pp. 836–840. Available at: 10.1038/s41592-023-01866-3.

Ströh, S., Hammerschmith, E.W., Tank, D.W., Seung, H.S. and Wanner, A.A. (2022) “In situ X-ray-assisted electron microscopy staining for large biological samples,” eLife, 11, p. e72147. Available at: 10.7554/elife.72147.

Strugnell, J., Jackson, J., Drummond, A.J. and Cooper, A. (2006) “Divergence time estimates for major cephalopod groups: evidence from multiple genes,” Cladistics, 22(1), pp. 89–96. Available at: 10.1111/j.1096-0031.2006.00086.x.

Styfhals, R., Zolotarov, G., Hulselmans, G., Spanier, K.I., Poovathingal, S., Elagoz, A.M., Winter, S.D., Deryckere, A., Rajewsky, N., Ponte, G., Fiorito, G., Aerts, S. and Seuntjens, E. (2022) “Cell type diversity in a developing octopus brain,” Nature Communications, 13(1), p. 7392. Available at: 10.1038/s41467-022-35198-1.

Sweeney, A.M., Marais, D.L.D., Ban, Y.-E.A. and Johnsen, S. (2007) “Evolution of graded refractive index in squid lenses,” Journal of The Royal Society Interface, 4(15), pp. 685–698. Available at: 10.1098/rsif.2006.0210.

Turchetti-Maia, A., Shomrat, T. and Hochner, B. (2017) “The Oxford Handbook of Invertebrate Neurobiology,” pp. 559–574. Available at: 10.1093/oxfordhb/9780190456757.013.29.

Vidal, E.A.G. and Salvador, B. (2019) “The Tentacular Strike Behavior in Squid: Functional Interdependency of Morphology and Predatory Behaviors During Ontogeny,” Frontiers in Physiology, 10, p. 1558. Available at: 10.3389/fphys.2019.01558.

Villar, F., Andre, L., Baker, R., Bohic, S., Silva, J.C. da, Guilloud, C., Hignette, O., Meyer, J., Pacureanu, A., Perez, M., Salome, M., Linden, P. van der, Yang, Y. and Cloetens, P. (2018) “Nanopositioning for the ESRF ID16A Nano-Imaging Beamline,” Synchrotron Radiation News, 31(5), pp. 9–14. Available at: 10.1080/08940886.2018.1506234.

Vo, N.T., Atwood, R.C. and Drakopoulos, M. (2018) “Superior techniques for eliminating ring artifacts in X-ray micro-tomography.,” Optics express, 26(22), pp. 28396–28412. Available at: 10.1364/oe.26.028396.

Wells, M.J. and Wells, J. (1957) “The Effect of Lesions to the Vertical and Optic Lobes on Tactile Discrimination in Octopus,” Journal of Experimental Biology, 34(3), pp. 378–393. Available at: 10.1242/jeb.34.3.378.

Williamson, R. and Budelmann, B.U. (1985) “The response of theOctopus angular acceleration receptor system to sinusoidal stimulation,” 156(3), pp. 403–412. Available at: 10.1007/bf00610733.

Wollesen, T. (2015) “Structure and Evolution of Invertebrate Nervous Systems,” pp. 222–240. Available at: 10.1093/acprof:oso/9780199682201.003.0021.

Woo, T., Liang, X., Evans, D.A., Fernandez, O., Kretschmer, F., Reiter, S. and Laurent, G. (2023) “The dynamics of pattern matching in camouflaging cuttlefish,” Nature, 619(7968), pp. 122–128. Available at: 10.1038/s41586-023-06259-2.

Xu, C.S., Hayworth, K.J., Lu, Z., Grob, P., Hassan, A.M., García-Cerdán, J.G., Niyogi, K.K., Nogales, E., Weinberg, R.J. and Hess, H.F. (2017) “Enhanced FIB-SEM systems for large-volume 3D imaging,” eLife, 6, p. e25916. Available at: 10.7554/elife.25916.

Yamamoto, M., Shimazaki, Y. and Shigeno, S. (2003) “Atlas of the Embryonic Brain in the Pygmy Squid, Idiosepius paradoxus,” Zoological Science, 20(2), pp. 163–179. Available at: 10.2108/zsj.20.163.

Young, J.Z. (1961) “LEARNING AND DISCRIMINATION IN THE OCTOPUS,” Biological Reviews, 36(1), pp. 32–95. Available at: 10.1111/j.1469-185x.1961.tb01432.x.

Young, J.Z. (1962) “The optic lobes of Octopus vulgaris,” Philosophical Transactions of the Royal Society of London. Series B, Biological Sciences, 245(718), pp. 19–58. Available at: 10.1098/rstb.1962.0005.

Young, J.Z. (1971) “The Anatomy of the Nervous System of Octopus Vulgaris,” Oxford University Press [Preprint].

Young, J.Z. (1974a) “The central nervous system of Loligo I. The optic lobe,” Philosophical Transactions of the Royal Society of London. B, Biological Sciences, 267(885), pp. 263–302. Available at: 10.1098/rstb.1974.0002.

Young, J.Z. (1974b) “The central nervous system of Loligo. I. The optic lobe,” Philos. Trans. R. Soc. Lond. B Biol. Sci., (267), pp. 263–302.

Young, J.Z. (1976) “The nervous system of Loligo. II. Subesophageal centers,” Philos. Trans. R. Soc. Lond. B Biol. Sci., (274), pp. 101–167.

Young, J.Z. (1977) “The nervous system of Loligo. III. Higher motor centers - the basal supraesophageal lobes,” Philos. Trans. R. Soc. Lond. B Biol. Sci., (276), pp. 351–398.

Young, J.Z. (1979) “The nervous system of Loligo. V. The vertical lobe complex,” Philos. Trans. R. Soc. Lond. B Biol. Sci., (285), pp. 311–354.

Yu, B., Weber, L., Pacureanu, A., Langer, M., Olivier, C., Cloetens, P. and Peyrin, F. (2018) “Evaluation of phase retrieval approaches in magnified X-ray phase nano computerized tomography applied to bone tissue,” Optics Express, 26(9), p. 11110. Available at: 10.1364/oe.26.011110.

