## Supplementary data for "Mapping the nervous system of the Idiosepius *hallami* pygmy squid: insights from whole-animal X-ray nanotomography imaging"

### Supplementary material

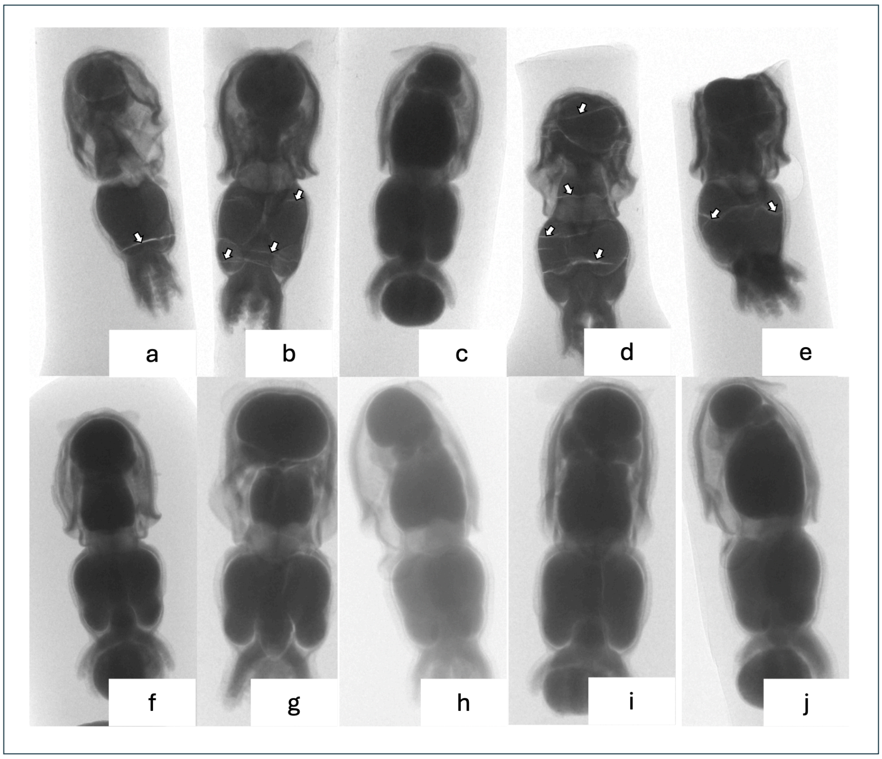


**Figure S1. Embed812 vs Durcupan microCTs. -** a-e) microCT projections of pygmy squid samples embedded in Embed812. Crack artefacts (white arrows) were visible(except in sample c). Sample C had too much visible yolk, which could place the pygmy squid in an earlier staging than other samples; f-j) microCT projections of pygmy squid samples embedded in Durcupan, no cracks are visible (Silva, 2024).
